# Enhanced inflammatory response mediated by parenchymal cells associates with resistance towards mTOR inhibition

**DOI:** 10.1101/2020.08.06.239426

**Authors:** Long Jiao, Roman Eickhoff, Sandra Jumpertz, Johanna Roth, Merve Erdem, Andreas Kroh, Hans Duimel, Carmen López-Iglesias, Maximilian Schmeding, Lara R. Heij, David Meierhofer, Ulf P. Neumann, Thorsten Cramer

## Abstract

Activation of the mTOR pathway is frequently found in cancer, but mTOR inhibitors have thus far failed to demonstrate significant antiproliferative efficacy in the majority of cancer types. Besides cancer cell-intrinsic resistance mechanisms, it is conceivable that mTOR inhibitors impact on non-malignant host cells in a manner that ultimately supports resistance of cancer cells. Against this background, we sought to analyze the functional consequences of mTOR inhibition in hepatocytes for the growth of metastatic colon cancer. To this end, we established a <u>l</u>iver <u>e</u>pithelial <u>c</u>ell (LEC)-specific knock-out (KO) of mTOR (mTOR^LEC^ mice). We used these mice to characterize the growth of colorectal liver metastases with and without partial hepatectomy to model different clinical settings. While the LEC-specific loss of mTOR remained without effect on metastasis growth in intact liver, partial liver resection resulted in the formation of larger metastases in mTOR^LEC^ mice compared to wildtype controls. This was accompanied by significantly enhanced inflammatory activity in LEC-specific mTOR KO livers after partial liver resection. Analysis of NF-κB target gene expression and immunohistochemistry of p65 displayed a significant activation of NF-κB in mTOR^LEC^ mice, suggesting a functional importance of this pathway for the observed inflammatory phenotype. Taken together, we show an unexpected acceleration of liver metastases upon deletion of mTOR in liver epithelial cells. Our results support the notion that non-malignant host cells can contribute to resistance against mTOR inhibitors and encourage to test if anti-inflammatory drugs are able to improve the efficacy of mTOR inhibitor for cancer therapy.

## Introduction

Colorectal cancer (CRC) ranks as the third most frequent malignant tumor worldwide and is accompanied by substantial cancer-associated morbidity and mortality (1). While the prognosis of patients with locally confined tumor growth has improved significantly in recent years (2), clinical care of metastasized CRC remains challenging (3). Due to continuous development of surgical techniques and establishment of interdisciplinary therapy planning, CRC patients with liver metastases can nowadays undergo treatment with curative intent (4). This is an impressive achievement and about 25% of patients survive five years or longer (4). Unfortunately, the majority of patients still develop tumor recurrence in the liver or other organs and will, ultimately, succumb to the disease (2). This demonstrates the urgent need to better understand the pathogenesis of both primary and secondary liver metastasis of colorectal cancer.

Our understanding of the molecular and cell biological mechanisms that govern the function of non-tumor cells for organ metastasis have greatly improved in recent years. The vast majority of publications have focussed on the role of immune cells, fibroblasts and endothelial cells of the tumor microenvironment for metastatic niche formation (5). Rather astonishingly, the function of hepatocytes, by far the most abundant cell type in the liver, for hepatic metastasis received a lot less attention. It has been reported over 30 years ago that specific lectins on hepatocytes are functionally important for liver metastasis via mediating adherence of tumor cells to the liver (6). Shortly thereafter, the group of Pnina Brodt showed paracrine stimulation of metastatic cell growth by IGF-1 secreted from hepatocytes (7). It was not until recently, however, that the mechanistic importance of hepatocytes for directing metastatic niche formation has received substantially more attention. A very elegant study directed by Gregory Beatty has shown that interleukin-6 (derived from the microenvironment of pancreas and colon cancer) stimulates hepatocytes to release serum amyloid A (and other factors) to induce accumulation of myeloid cells and activation of hepatic stellate cells, thereby supporting metastatic seeding and outgrowth (8). With an equally impressive experimental approach, Georg Halder and colleagues were able to show that both primary and secondary liver tumors resulted in activation of the Hippo downstream effectors YAP and TAZ in hepatocytes and that experimental hyperactivation of YAP in hepatocytes surrounding melanoma-derived liver metastasis inhibited tumor growth (9). These studies convincingly demonstrate that hepatocytes are able to support as well as to constrain metastatic progression in the liver and underscore the need to better understand the functionally involved pathways.

Metabolic reprogramming is a hallmark of both primary tumor formation and metastasis (10). Besides cellular growth, antioxidant defence and therapy resistance, tumor-specific metabolic alterations are of pivotal importance for various stages of metastasis (11). While cancer metabolism was for the longest time thought to be primarily determined by cell-intrinsic factors, e.g. genetic changes, we know today that the tumor microenvironment (TME) is able to profoundly modify the metabolic activity of cancer cells (12). Breast cancer cells, for example, display significant differences in their use of anaplerotic pathways between primary tumor and lung metastases, suggesting a causal involvement of the TME (13). With respect to colon cancer it was shown that creatine in the extracellular space of the liver is used by cancer cells for ATP generation, ultimately facilitating survival and growth of hepatic metastasis (14). These fascinating results led us to hypothesize that intervening with the metabolism of hepatocytes could inhibit the survival of colon cancer metastases in the liver, potentially representing a novel therapeutic approach. To test our hypothesis, we evaluated the consequences of a functional loss of the mTOR (mechanistic target of rapamycin) pathway in hepatocytes. The highly conserved serine/threonine kinase mTOR translates nutrient availability and growth factor signaling into different cellular responses (15). Under conditions of abundant nutrients (e.g. glucose and amino acids), growth and proliferation of cells is encouraged by mTOR (16). Conversely, when nutrients are scarce, mTOR signaling is inhibited, resulting in a reduction of anabolic processes (e.g. protein and lipid synthesis), induction of autophagy and, ultimately, cessation of cellular proliferation (17). mTOR consists of two functionally and molecularly distinct complexes: mTOR complex 1 (mTORC1) is activated by nutrients and growth factor signaling and inhibited by rapamycin while mTORC2 is mainly activated by growth factors and insensitive towards acute rapamycin treatment (15). Taken together, mTOR is considered to be a central signaling relay for the orchestration of cellular metabolism with environmental conditions.

## Materials and Methods

### Animal experiments

All experiments were done in accordance with the German legislation governing animal studies and the Guide for the Care and Use of Laboratory Animals (National Institutes of Health (NIH), Publication No 85-23, revised 2011). The experiments were approved by the Governmental Animal Care and Use Committee (Landesamt für Natur Umwelt und Verbraucherschutz (LANUV) Nordrhein-Westfalen, Recklinghausen, Germany; reference number: 84-02.04.2015.A126 and 81-02.04.2017.A398). <u>L</u>iver <u>e</u>pithelial <u>c</u>ell-specific mTOR knock-out (KO) mice (termed mTOR^LEC^) were generated via the Cre/loxP (18) system with floxed mTOR mice (19) and albumin-Cre mice (20). 8-12 weeks old mice were used for experiments. Mice were housed in the Institute for Laboratory Animal Science (ILAS) at the University Hospital of the RWTH Aachen University. Mice were given ad libitum access to water and chow. Gastight room under strict 12h light/dark cycles (day: 7:00 am-7:00 pm), H_2_O_2_ gas-disinfection, cage bedding, 22 ± 1°C room temperature, 50 ± 10% relative humidity and specific pathogen free (SPF) conditions were applied.

### Antibodies

Phospho-Akt (Ser473) (Cell Signaling Technology (CST), #4060), β-actin (Sigma-Aldrich #A5441), Caspase-3 (Abcam, #ab4051), CD45 (Thermo Fisher Scientific, #14-0451-82), F4/80 (Thermo Fisher Scientific, #14-4801-82), Ki-67 (CST, #12202), Ly-6G (BioLegend, #127601), LC3B (Sigma-Aldrich, #L7543), mTOR (CST, #2983), p62 (SQSTM1) (MBL International, #PM045), p65 (CST, #8242), p-p70-S6 Kinase (CST, #9234), p-Histone H3 (CST, #9701), RIP3 (ProSci, #2283).

### Cell lines and primary hepatocytes

The established murine colon carcinoma cell line MC38 (21) was maintained in Dulbecco’s Modified Eagle’s Medium Ham’s Nutrient Mixture F-12 (DMEM, Sigma-Aldrich, D8437) with 10% fetal bovine serum (Gibco). Cells were used between passages 10 and 25 at approximately 70-90% confluence. Primary murine hepatocytes were isolated from livers of WT and mTOR^LEC^ mice at 8-12 weeks of age and maintained in DMEM (Gibco) with 10% fetal bovine serum as previously reported (22). Cells were cultured in a humidified incubator with 5% CO_2_ at 37°C.

### Murine model of colorectal cancer liver metastasis

All surgical procedures were performed under general anaesthesia. Analgesia was ensured by Buprenorphine (0.05 mg/kg BW) 30 minutes before skin incision and in 8h intervals for 3 days after the operation. Anesthesia was performed with an isoflurane-oxygen mixture (induction: 3-4% isoflurane, maintenance: 1-1.5% isoflurane, oxygen flow 2l/min), and mice were fixed in a supine position on a heating pad. The abdominal cavity was exposed through a 3-cm midline laparotomy and an abdominal wall retractor system was placed. The spleen was exposed on a gauze and 7.5 × 10^4^ MC38 cells in 250μl HBSS (Hank’s balance salt solution, Gibco) were injected into the central part of the spleen within 15 seconds, and the needle was maintained in the spleen for 2 minutes. Afterwards, the spleen was removed and the abdominal wall was closed by suturing the muscle layer followed by the skin. After 14 days, mice were anesthetized as described before and the abdominal cavity was re-opened. The left-anterior and right-anterior segment of the liver including the gallbladder were resected by haemostatic clip ligation according to published literature (23). After establishing haemostasis, the abdominal cavity was closed and analgesia was continued for 3 days. All mice were sacrificed 4 weeks after the initial operation, and the number of metastatic nodules was evaluated.

### Isolation of genomic DNA and total RNA and quantitative PCR analysis

Genomic DNA (gDNA) was isolated from mouse ear punches and hepatocytes with the NucleoSpin Tissue Kit (Macherey-Nagel, Germany). RNA was isolated from mouse liver with peqGOLD RNAPure reagent (VWR, USA) and QIAshredder spin columns (Qiagen) according to the manufacturer’s instructions. RNA yield and purity were determined spectrophotometrically. RNA integrity was checked by MOPS buffered denaturizing RNA gel electrophoresis and only samples with a 28S/18S ratio of at least 1.8 were included in the qPCR study. First strand cDNA of 1 μg total RNA per reaction was synthesized with an oligo (dT) primer and a SuperScript™ First Strand Synthesis System (Invitrogen). Quantitative real-time PCR analysis was conducted on an ABI 7500 Real-Time PCR system using PowerSYBR® Green PCR Master Mix (Thermo Fisher Scientific, Waltham, USA). Primer-specific annealing temperatures were pre-evaluated prior to the study to optimize PCR conditions. Primer specificity was checked by melt curve analyses and TAE-buffered DNA agarose gel electrophoresis of obtained PCR products. Amplification efficiency was calculated with LinRegPCR 2016.0 (Heart Failure Research Center, Amsterdam, The Netherlands) and used for efficiency corrections (24). Inter-Run calibrators were used if necessary to correct for inter-run variations. Relative fold-changes of target gene expression were calculated with qbase+ 3.0 (Biogazelle, Zwijnaarde, Belgium) that automatically applies the above mentioned corrections to the ΔΔCq method (25). Primer sequences are available upon request.

### Protein Isolation and Western Blot Analysis

Hepatocytes or liver tissue samples were lysed in RIPA lysis buffer (10mmol/L Tris-HCl pH7.5, 150mmol NaCl, 0.25% SDS, 10mg/ml Natriumdeoxycholat, 1% Nonidet P40, 1mmol/L Na_3_VO_4_, 1mmol/L DTT, 2mmol/L PMSF, 10mmol/L NaF, 2μmol/L Leupeptin and 4.4×10^−4^ TIU/mg Aprotinin). The lysate was sonicated and clarified by centrifugation (12.000g, 10 minutes, 4°C), snap frozen in liquid nitrogen and stored at −80°C until assayed. Total protein samples (30μg) were heated at 95°C for 5 min in sample buffer, subjected to polyacrylamide gel electrophoresis (PAGE) and electro-transferred onto nitrocellulose membranes (GE Healthcare, Germany). Blocking with 5% non-fat dry milk or bovine serum albumin (BSA, Carl Roth, Germany) was done for 1h at room temperature (RT). Blocking buffer was removed and the membranes were incubated with the primary antibody overnight at 4°C. After washing three times for 5 min in TBST (TBS buffer with 0.1% v/v Tween-20), the secondary antibody was applied for 1h at RT. Thereafter, the membrane was washed thrice for 5 min with TBST, incubated for 1 min ECL (Perkin Elmer, USA) and the signal was determined with an INTAS ECL Chemocam imager (INTAS Science Imaging, Germany).

### Immunohistochemistry

All Immunohistochemistry experiments were performed on formalin-fixed, paraffin-embedded tissues with heat-mediated antigen retrieval (10 min at 110°C in Dako Target Retrieval Solution in the Decloaking Chamber (Biocare Medical, USA)). After washing twice with TBST, the slides were blocked with blocking buffer (ZytoChem Plus AP kit, Zytomed System, Germany) for 5 min, followed by Avidin and Biotin blocking for 15 min (Avidin/Biotin Blocking Kit, Vector, Germany). Slides were incubated with primary antibody overnight at 4°C, followed by two washing steps and application of the secondary antibody for 1h at RT. After repeated washing with TBST, subjection to streptavidin-AP conjugate (ZytoChem Plus AP kit) was done for 15 min at RT and the reaction was developed with AP red solution (Permanent AP Red kit, Zytomed System). The reaction was stopped by washing the slides briefly in demineralized water (dH_2_O), and nuclei were briefly stained with haematoxylin.

### Oil red O staining

Cryosections were done at 10 μm and immediately fixed in ice-cold formalin for 10 min. After washing with PBS for 5 min, sections were incubated in 60% 2-propanol for 10 seconds. Oil red O staining solution was applied for 15 min, followed by running tap water for 5 min and nuclear were briefly stained with haematoxylin for 30 seconds. The reaction was stopped in dH_2_O and slides were mounted with glycerol gelatin (Glycergel, Dako) and coverslipped.

### Electron microscopy

Liver tissue of WT and mTOR^LEC^ mice was fixed with 1.5% glutaraldehyde in 0.067M cacodylate buffer at pH 7.4 containing 1% sucrose and kept in the fixative for 24h at 4°C. Then, tissue samples were washed with 0.1M cacodylate buffer with sucrose and post-fixed with 1% osmium tetroxide in the same buffer containing 1.5% potassium ferricyanide for 1h at 4°C.

Then, samples were washed in 0.1 M cacodylate followed by dehydration in ethanol, infiltration with Epon resin for 2d, embedding in the same resin and polymerisation at 60°C for 48h. Ultrathin sections were obtained using a Leica Ultracut UCT ultramicrotome and mounting on Formvar-coated copper grids. These were staining with 2% uranyl acetate in water and lead citrate. Finally, sections were analyzed in a Tecnai T12 electron microscope equipped with an Eagle 4kx4k CCD camera (Thermo Fisher Scientific, The Netherlands).

### Proteomics sample preparation with label-free quantification (LFQ)

About 10 mg frozen liver tissue per sample was homogenized under denaturing conditions with a FastPrep (three times for 60 s, 4 m × s-1) in 1 mL of a buffer containing 4% SDS, 0.1 M DTT, 0.1 M Tris pH 7.8, followed by sonication for 1 min, boiled at 95°C for 5 min in a thermal shaker and centrifuged at 15,000 g for 5 min. The supernatant was transferred into new protein low binding tube (Eppendorf, Germany). Protein precipitation was achieved by adding four times excess volume of ice-cold acetone at −20°C overnight. Pelleting at maximum speed at 4 °C was followed by three washes with acetone and samples were dried in a vacuum concentrator. Lyophilized proteins were dissolved in 6 M guanidinium chloride (GdmCl), 5 mM tris(2-carboxyethyl)phosphine, 20 mM chloroacetamide and 50 mM Tris-HCl pH 8.5. Samples were boiled for 5 min at 95 °C and sonicated for 15 min in a water sonicator. About 150 μg protein per sample (~10 μL) were diluted 1:10 with nine times volume of 10% acetonitrile and 25 mM Tris pH 8.5, followed by a trypsin digestion (1:50) at 37 °C overnight. Subsequent peptides were purified with C18 columns and further fractionated by strong cation exchange (SCX) chromatography. Desalted peptides were reconstituted in 0.1% formic acid in water and further separated into four fractions by strong cation exchange chromatography (SCX, 3M Purification, Meriden, CT). Eluates were first dried in a SpeedVac, then dissolved in 10 μl 5% acetonitrile and 2% formic acid in water, briefly vortexed, and sonicated in a water bath for 30 seconds prior injection to nano-LC-MS/MS. 5 μg of each SCX fraction and a non-fractioned sample were used for proteome profiling.

### LC-MS/MS instrument settings for shotgun proteome profiling and data analysis

LC-MS/MS was carried out by nanoflow reverse-phase liquid chromatography (Dionex Ultimate 3000, Thermo Scientific) coupled online to a Q-Exactive Plus Orbitrap mass spectrometer (Thermo Scientific). Briefly, the LC separation was performed using a PicoFrit analytical column (75 μm ID × 55 cm long, 15 μm Tip ID; New Objectives, Woburn, MA) in-house packed with 2.1-μm C18 resin (Reprosil-AQ Pur, Dr. Maisch, Ammerbuch, Germany). Peptides were eluted using a non-linear gradient from 3.8 to 50% solvent B over 173 min at a flow rate of 266 nl/min (solvent A: 99.9% water, 0.1% formic acid; solvent B: 79.9% acetonitrile, 20% water, 0.1% formic acid). 3.5 kilovolts were applied for nanoelectrospray ionization. A cycle of one full FT scan mass spectrum (300−1,750 m/z, resolution of 70,000 at m/z 200, AGC target 1 × 10^6^) was followed by 12 data-dependent MS/MS scans (200-2,000 m/z, resolution of 35,000, AGC target 5 × 10^5^, isolation window 2 m/z) with normalized collision energy of 25 eV. Target ions already selected for MS/MS were dynamically excluded for 30 seconds. In addition, only peptide charge states between two to eight were allowed. Raw MS data were processed with MaxQuant software (v1.6.10.43) and searched against the murine proteome database UniProtKB with 55,153 entries, released in 08/2019. Parameters of MaxQuant database searching were: A false discovery rate (FDR) of 0.01 for proteins and peptides, a minimum peptide length of 7 amino acids, a first search mass tolerance for peptides of 20 ppm and a main search tolerance of 4.5 ppm, and using the function “match between runs”. A maximum of two missed cleavages was allowed for the tryptic digest. Cysteine carbamidomethylation was set as fixed modification, while N-terminal acetylation and methionine oxidation were set as variable modifications. Contaminants, as well as proteins identified by site modification and proteins derived from the reversed part of the decoy database, were strictly excluded from further analysis. The mass spectrometry data have been deposited to the ProteomeXchange Consortium (http://proteomecentral.proteomexchange.org) via the PRIDE partner repository (26) with the dataset identifier PXD019998.

### Experimental design, statistical rationale, pathway and data analyses

The correlation analysis of biological replicates and the calculation of significantly different proteins was done with Perseus (v1.6.10.43). LFQ intensities, originating from at least two different peptides per protein group, were transformed by log2. Only protein groups with valid values within compared experiments were used for further data evaluation. Statistical analysis was done by a two-sample t-test with Benjamini-Hochberg (BH, FDR of 0.05) correction for multiple testing. For comprehensive proteome data analyses, we applied gene set enrichment analysis (GSEA, v3.0) (27) in order to see, if *a priori* defined sets of proteins show statistically significant, concordant differences between knockouts and controls. We used GSEA standard settings, except that the minimum size exclusion was set to 5 and Reactome v7.0 and KEGG v7.0 were used as gene set databases. The cut-off for significantly regulated pathways was set to be ≤ 0.05 p-value and ≤ 0.25 FDR.

## Results

### Basic characterization of liver epithelial cell-specific mTOR knockout mice

To investigate the functional role of mTOR in hepatocytes for liver metastasis, we generated <u>l</u>iver <u>e</u>pithelial <u>c</u>ell-specific mTOR knockout mice (mTOR^LEC^) via the Cre/loxP system (18). Quantitative PCR with genomic DNA isolated from primary hepatocytes displayed a deletion efficiency of roughly 50% (Figure S1A). Immunoblot analysis of primary hepatocytes isolated from mTOR^LEC^ mice revealed a strong reduction of the mTOR protein, while whole liver lysates failed to show a significant difference between WT and mTOR^LEC^ mice (Figure S1B). To analyze the functional effects of the mTOR loss, established downstream target proteins of mTOR were characterized via immunoblot. As can be seen in figure S1C, both basal and stimulated phosphorylation of p70/S6K is clearly reduced in mTOR-deficient hepatocytes. Insulin-stimulated activation of Akt was enhanced in KO hepatocytes, confirming earlier reports (28, 29). Overall, these results demonstrated sufficient inhibition of the mTOR signaling pathway in mTOR^LEC^ mice. To characterize the effect of the LEC-specific mTOR loss on the global proteome, label-free mass spectrometry was performed. Pearson correlation was used to compare the proteome profiles of all five biological replicates of WT and mTOR^LEC^ mice. The Pearson correlation coefficients were highly similar, ranging from 0.926 to 0.99 in WT and KO liver tissue (Figure S2A), suggesting a very good quality of the proteome data sets. A volcano plot visualized proteins with significantly different expression between the genotypes (Figure S2B, Table 1). With the help of the STRING (Search Tool for the Retrieval of Interacting Genes/Proteins) software, the network of molecular interactions among downregulated proteins (at least one standard deviation from the mean) was visualized (Figure S3). The most striking and significantly downregulated pathways within this network were RNA binding (FDR: 3.62e-09, GO) and the ribosome (FDR: 1.03e-05, KEGG), both known to be under control of the mTOR pathway, further supporting functional mTOR inactivation (30). Taken together, the LEC-specific loss of mTOR had only modest effects on the proteome profile, which can be explained either by insufficient KO efficiency or by activation of compensatory pathways. Next, we sought to evaluate the impact of the LEC-specific mTOR loss on liver histology. Various routine histochemical stainings (H&E, Sirius red and Oil red O) did not show differences between WT and mTOR^LEC^ mice (Figure 1). Likewise, hepatocellular proliferation (judged by Ki-67 nuclear positivity, figure S4) and apoptosis (using cleaved caspase-3, figure S4) was not different between the genotypes. Given the importance of hepatocytes for hepatic macrophages (31), we sought to characterize the effect of the LEC-specific mTOR loss on the abundance of inflammatory cells in the liver. While the number of total leukocytes (visualized by CD45 immunohistochemistry) was unchanged, significantly higher numbers of F4/80-positive macrophages were detected in LEC-specific mTOR knockout mice (Figure 2). As treatment of myeloid cells with rapamycin, a well-established inhibitor of mTORC1, promotes the secretion of pro-inflammatory factors (e.g. tumor necrosis factor-α (TNF-α) and interleukin-6 (IL-6) (32–34), we determined the expression of genes known to impact on the recruitment of inflammatory cells. As can be seen in figure 3, interleukin-1β *(Il1b)*, TNF-α (*Tnfa)*, Chemokine (C-C motif) ligand 2 *(Ccl2)*, macrophage migration inhibitory factor *(Mif)*, transforming growth factor-β1 *(Tgfb1)*, interleukin-6 (*Il6)* and the anti-inflammatory cytokine interleukin-10 *(II10)* were expressed at similar levels in the livers of WT and mTOR^LEC^ mice. Autophagy, a catabolic process affecting organelle degradation and protein turnover (35), is of central importance for liver biology and inhibited by mTOR (36, 37), suggesting that autophagy might be enhanced in mTOR^LEC^ mice. However, analysis of autophagy based on the detection of LC3B processing, p62 protein expression and electron microscopy-based visualization of autophagosomes could not detect significant differences between the livers of WT and mTOR^LEC^ mice (Figure S5). In summary, with the exception of enhanced macrophage abundance in KO livers, the LEC-specific functional inactivation of mTOR remained without significant effects on the majority of basic biological and histological traits.

**Table 1.**
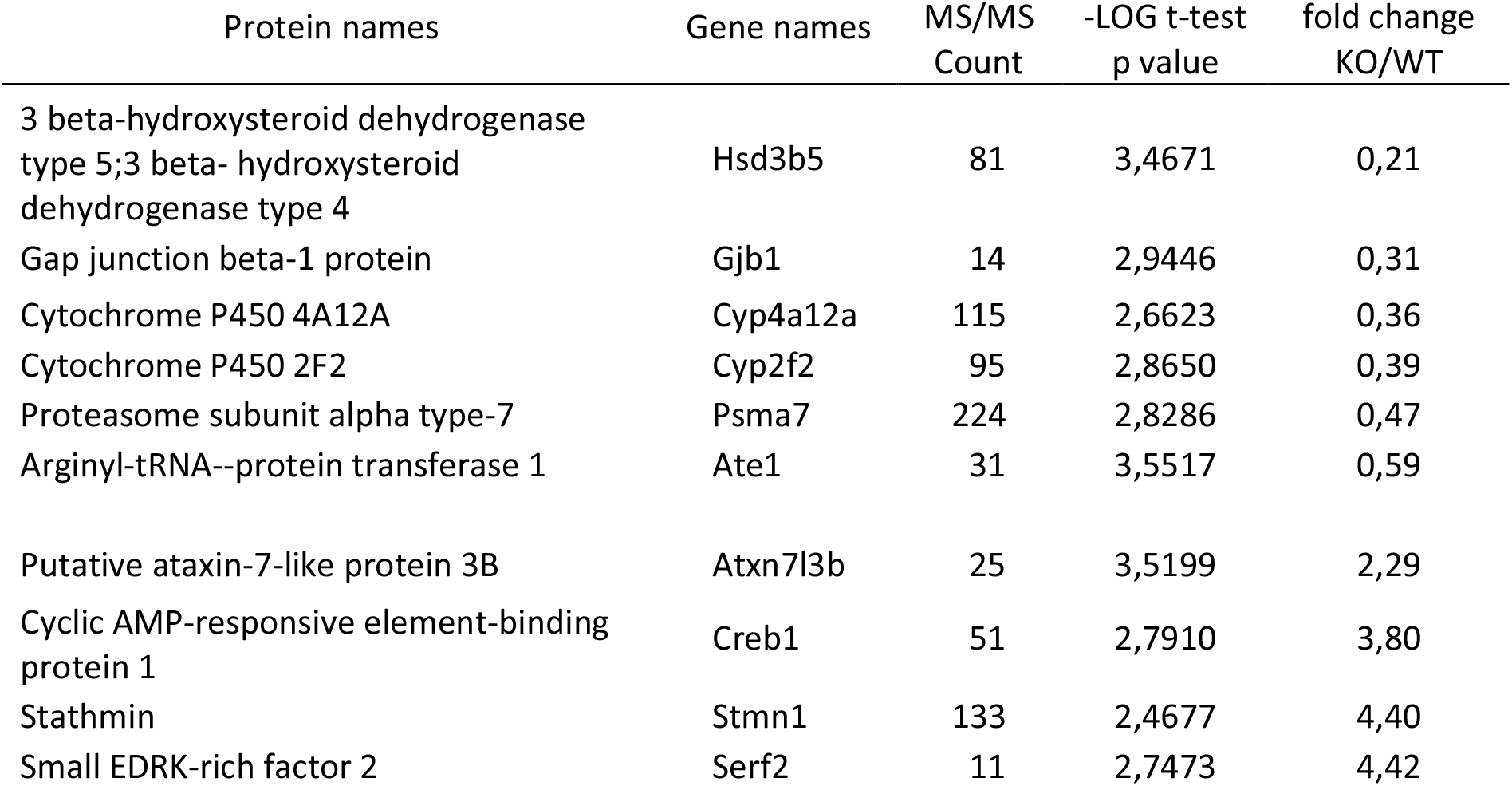
List of significantly altered proteins in mTOR^LEC^/WT liver tissue. Posterior error probability (PEP), MS2 spectral count frequency (MS/MS Count), Benjamini-Hochberg corrected p values and fold changes are shown.

| Protein names | Gene names | MS/MS<br>Count | -LOG t-test<br>p value | fold change<br>KO/WT |
| --- | --- | --- | --- | --- |
| 3 beta-hydroxysteroid dehydrogenase<br>type 5;3 beta- hydroxysteroid<br>dehydrogenase type 4 | Hsd3b5 | 81 | 3,4671 | 0,21 |
| Gap junction beta-1 protein | Gjb1 | 14 | 2,9446 | 0,31 |
| Cytochrome P450 4A12A | Cyp4a12a | 115 | 2,6623 | 0,36 |
| Cytochrome P450 2F2 | Cyp2f2 | 95 | 2,8650 | 0,39 |
| Proteasome subunit alpha type-7 | Psma7 | 224 | 2,8286 | 0,47 |
| Arginyl-tRNA--protein transferase 1 | Ate1 | 31 | 3,5517 | 0,59 |
| Putative ataxin-7-like protein 3B | Atxn7l3b | 25 | 3,5199 | 2,29 |
| Cyclic AMP-responsive element-binding<br>protein 1 | Creb1 | 51 | 2,7910 | 3,80 |
| Stathmin | Stmn1 | 133 | 2,4677 | 4,40 |
| Small EDRK-rich factor 2 | Serf2 | 11 | 2,7473 | 4,42 |

**Figure 1.**
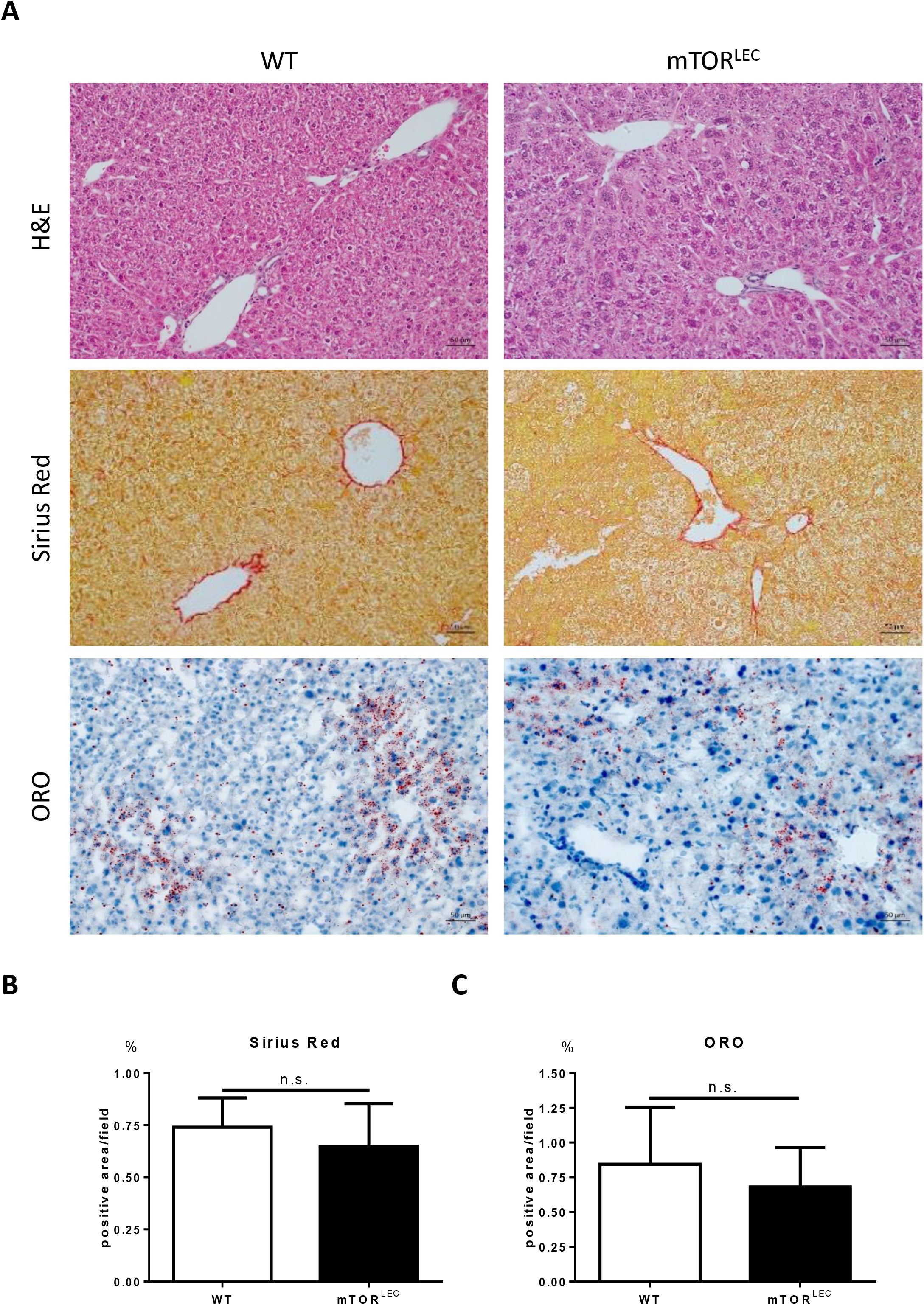
Basic histopathological traits are not affected by the LEC-specific mTOR loss. **A**, Liver sections from WT and mTOR^LEC^ mice were stained with H&E, Sirius Red and Oil red O (scale bar = 50 μm). **B,** Positive areas were quantified with ImageJ software and shown as bar graphs. Data are represented as mean + SEM by unpaired two-sided Student’s *t* test.

**Figure 2.**
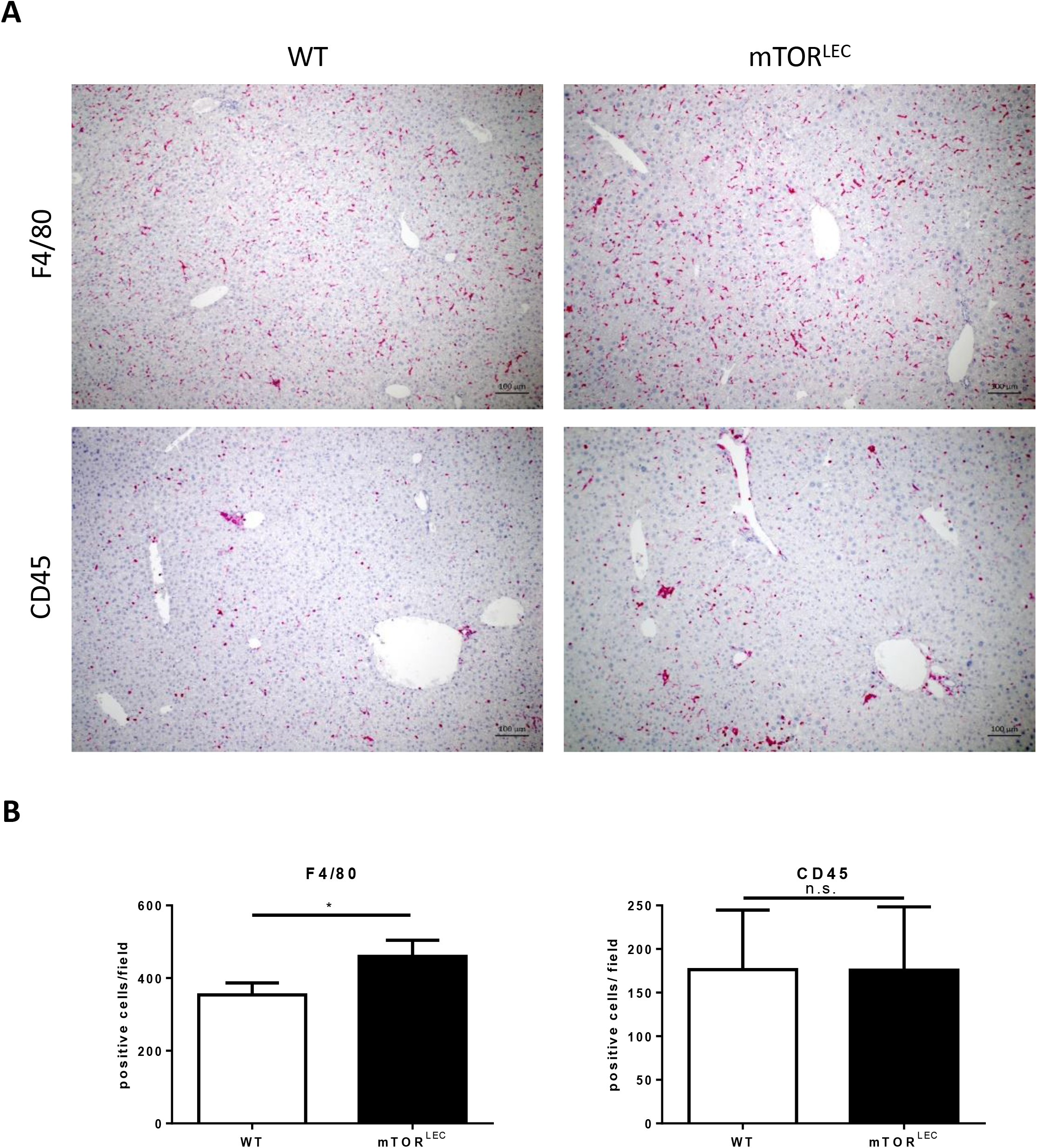
LEC-specific mTOR deletion results in elevated numbers of F4/80+ macrophages in the liver. **A**, Immunohistochemistry against F4/80 and CD45 of liver sections from WT and mTOR^LEC^ mice (scale bar = 100 μm). **B**, Positive cells were quantified with ImageJ software and shown as bar graphs. Data are represented as mean + SEM by unpaired two-sided Student’s *t* test, \**p* < 0.05.

**Figure 3.**
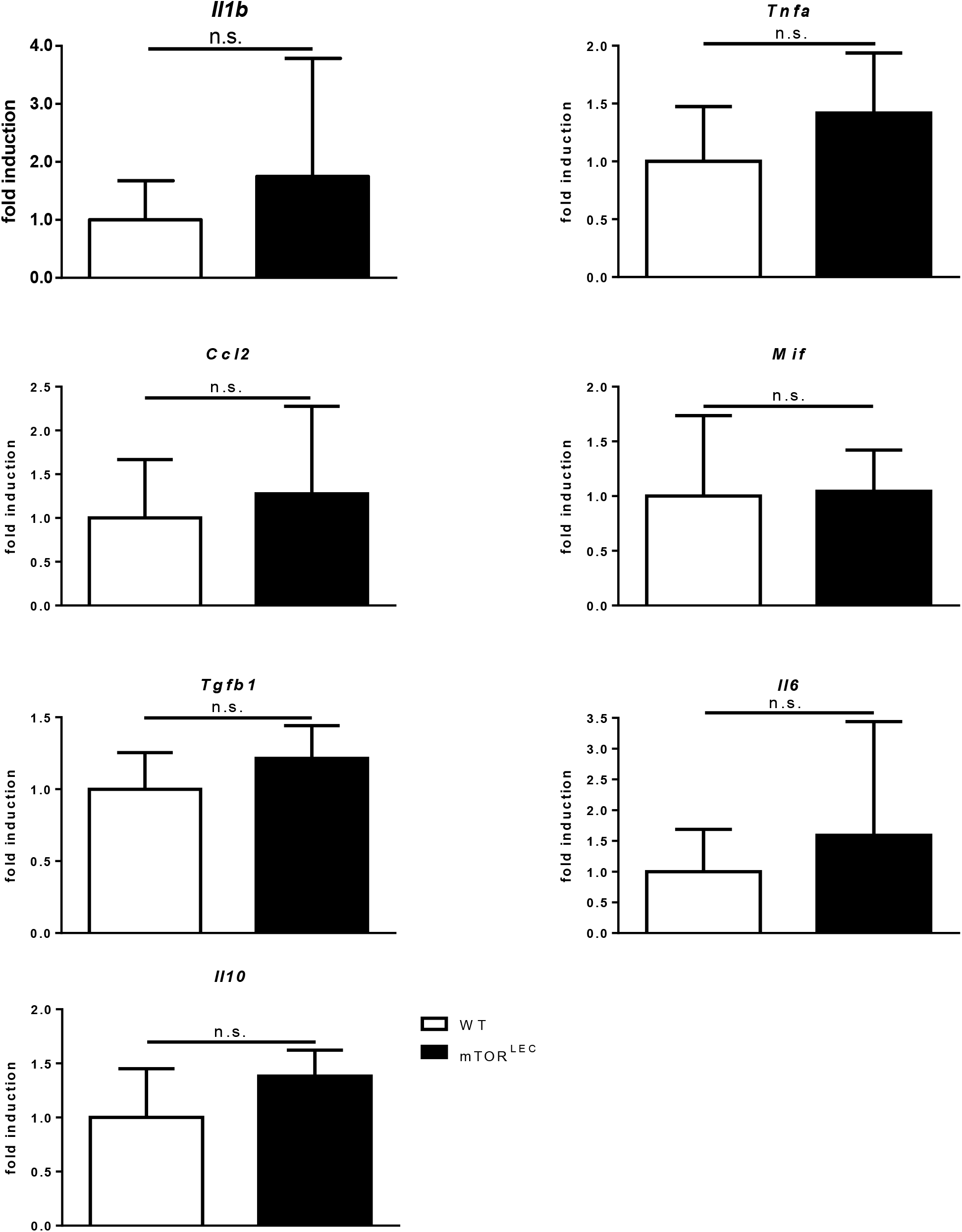
The expression of genes involved in the inflammatory response in the liver is not affected by the LEC-specific loss of mTOR. Gene expression was determined via quantitative real-time PCR, data were analyzed with qBase plus and the results are shown as bar graphs (KO relative to WT). Neither Interleukin-1β (*Il1b*), Tumor necrosis factor-α (*Tnfa*), Chemokine (C-C motif) ligand 2 (*Ccl2*), Macrophage migration inhibitory factor (*Mif*), Transforming growth factor-β1(*Tgfb1*), Interleukin-6 (*II6*) nor Interleukin-10 (Il10) are significantly different between the genotypes. Data are represented as mean + SEM by unpaired two-sided Student’s *t* test.

### Liver metastasis of colon cancer cells is not affected by LEC-specific loss of mTOR

Liver metastases of colon cancer in mice were established according to an established protocol (38). Injection of the established murine colon cancer cell line MC38 (21) into the spleen (with subsequent splenectomy) resulted in the formation of visible liver metastasis after 28 days (Figure 4A). Quantification of the metastatic load revealed no difference between WT and mTOR^LEC^ mice (Figure 4A). In line with this result, the relative number of proliferating cells both in the benign liver adjacent to the metastases and in the tumor nodules themselves were similar between the genotypes (Figure S6). Immunohistochemistry was applied to characterize the inflammatory reaction of the liver surrounding the metastatic nodules. Total leukocytes (CD45) and neutrophils (Ly6G) displayed no significant difference when WT and mTOR^LEC^ mice were compared (Figure 4B). Of note, the abundance of F4/80-positive macrophages was significantly higher upon LEC-specific loss of mTOR (Figure 4B), reminiscent of the findings in un-challenged mice (Figure 2).

**Figure 4.**
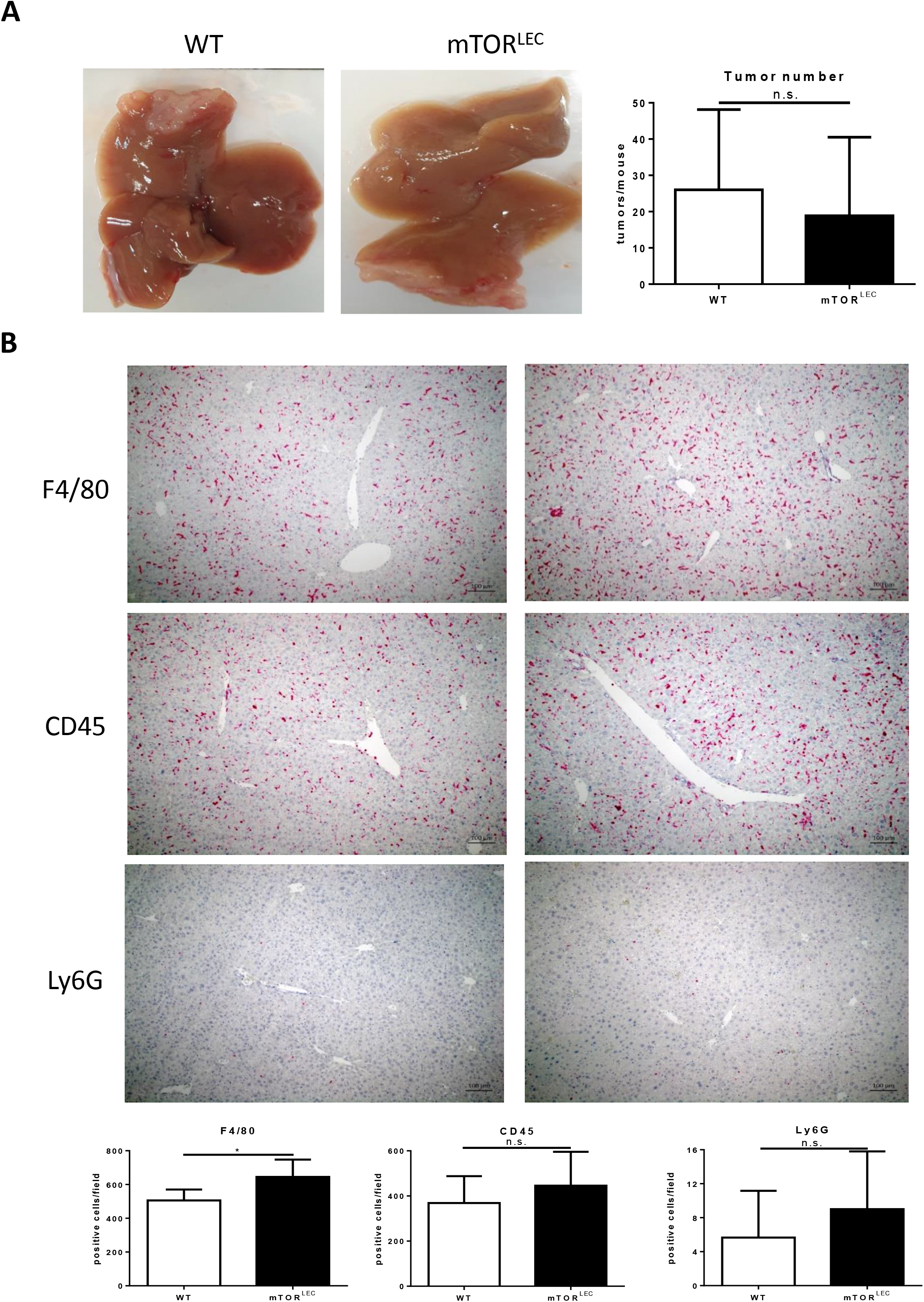
Loss of mTOR in liver epithelial cells does not affect murine colon cancer liver metastasis. **A**, Gross appearance of livers with MC38 metastases. Tumor number from both WT and mTOR^LEC^ mice were quantified with ImageJ software and shown as bar graphs. **B**, Abundance of leukocytes (CD45), F4/80+ macrophages and neutrophils (Ly6G) was investigated with immunohistochemistry. Positive cells were quantified with ImageJ and results are shown as bar graphs. Data are represented as mean + SEM by unpaired two-sided Student’s *t* test, \**p* < 0.05.

### Loss of mTOR in LECs enhances liver metastasis after partial hepatectomy

To be able to mirror the clinical situation of surgical resection of liver metastases, we performed partial liver resection 14 days after injection of the murine colon cancer cell line into the spleen (Figure S7). This resulted in a clear acceleration of metastasis progression in the mTOR^LEC^ mice (Figure 5A). Analysis of proliferation of metastasis-adjacent liver cells and tumor cells again showed no difference between WT and mTOR^LEC^ mice (Figure S8). Of note, the inflammatory activity in the benign liver surrounding the metastatic nodules was significantly enhanced in mice harbouring a LEC-specific mTOR deletion. All examined cell types (total leukocytes, F4/80-positive macrophages and neutrophils) were found at significantly higher levels in mTOR^LEC^ compared to WT mice (Figure 5B). We sought to better understand the underlying mechanism and decided to analyze NF-kB activity as this pathway is both crucial for the inflammatory response (39) and has been shown to functionally interact with mTOR (40, 41). Immunohistochemistry-based localization of p65 demonstrated nuclear positivity in hepatocytes exclusively in mTOR^LEC^ mice (Figure 6A). Furthermore, various established NF-kB target genes with pro-inflammatory functions (*Il1b, Tnfa, Ccl2, Ccl5, Cxcl1, Cxcl2 and Cxcl5*) were expressed at significantly higher levels in the livers of LEC-specific mTOR KO mice (Figure 6B). These results suggested that the loss of mTOR in liver parenchymal cells aggravated the growth of colon cancer liver metastases via NF-kB-mediated inflammation.

**Figure 5.**
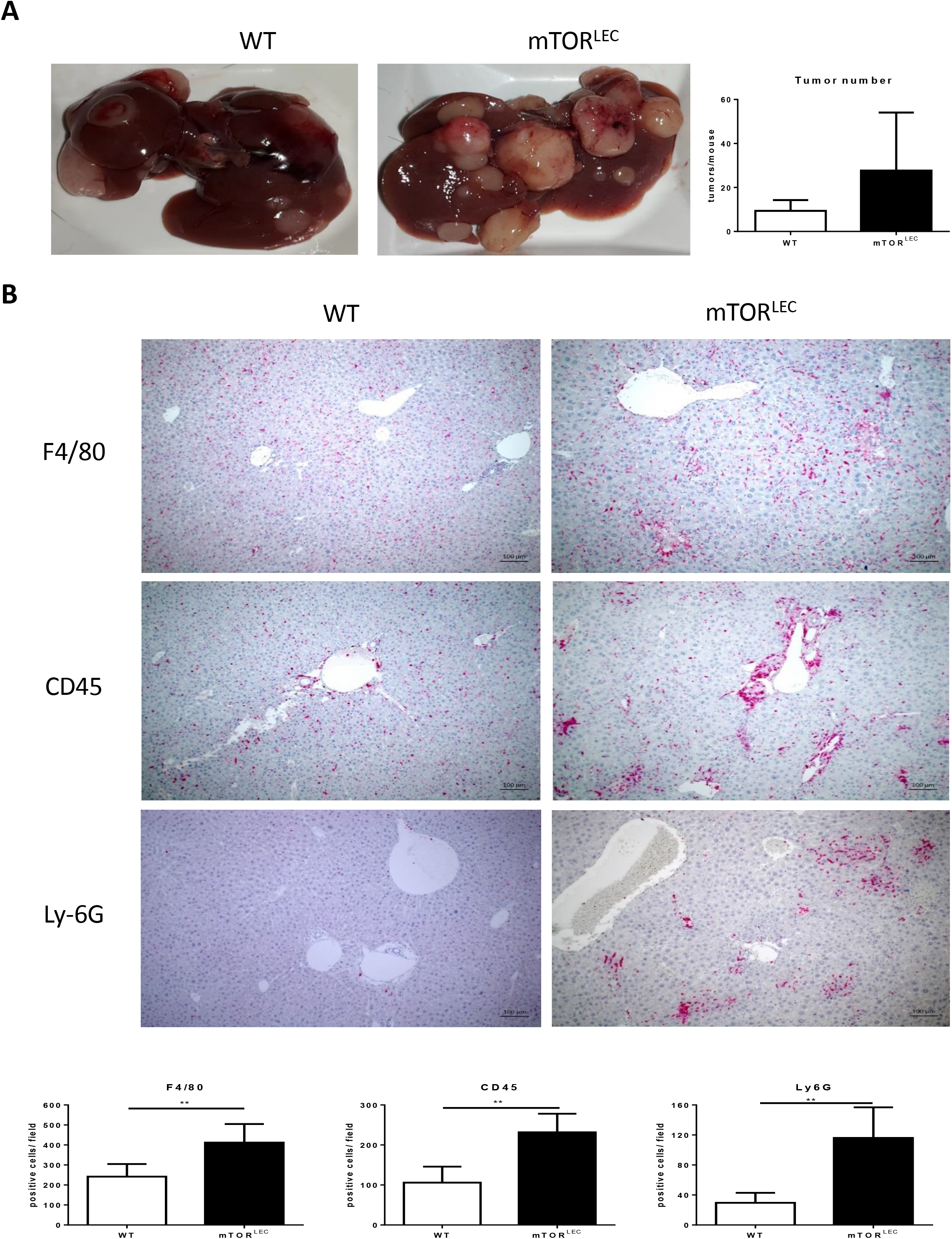
Enhanced metastatic load and inflammation after partial hepatectomy upon LEC-specific mTOR deletion. **A**, Gross appearance of livers with MC38 metastases after partial hepatectomy. Tumor number from both WT and mTOR^LEC^ mice were quantified with ImageJ software and shown as bar graphs. **B**, Abundance of leukocytes (CD45), F4/80+ macrophages and neutrophils (Ly6G) was investigated with immunohistochemistry. Positive cells were quantified with ImageJ and results are shown as bar graphs. Data are represented as mean + SEM by unpaired two-sided Student’s *t* test, \**p* < 0.05, \*\**p* <0.01.

**Figure 6.**
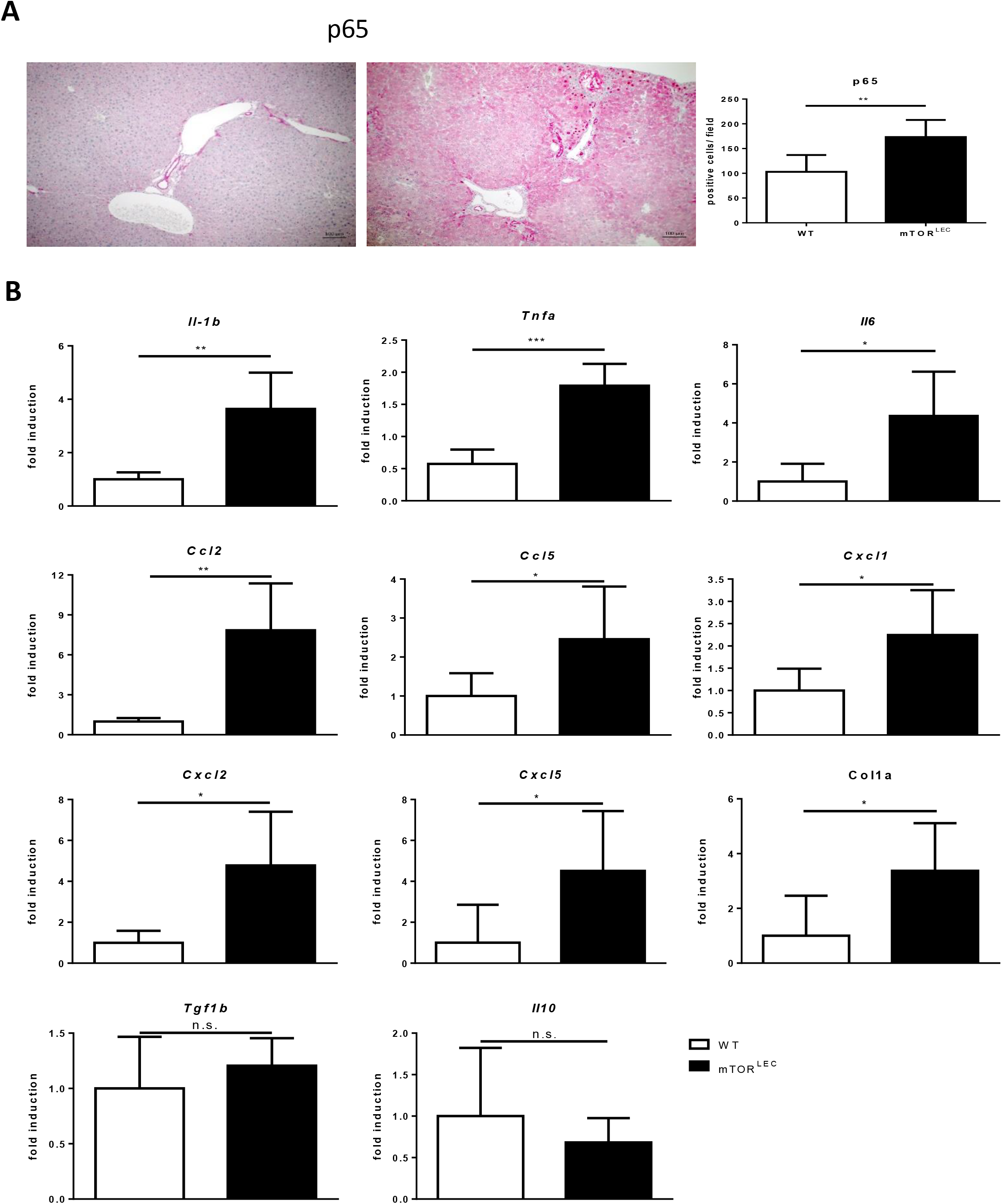
Elevated activity of the canonical NF-kB pathway in liver samples of mTOR^LEC^ mice after partial hepatectomy. Gene expression of established NF-kB target genes was determined via quantitative real-time PCR, data were analyzed with qBase plus and the results are shown as bar graphs (KO relative to WT). Interleukin-1β (*Il1b*), Tumor necrosis factor-α (*Tnfa*), Chemokine (C-C motif) ligand 2 (*Ccl2*), Chemokine(C-C motif) ligand 5 (*Ccl5*), Chemokine (C-X-C motif) ligand 1(*Cxcl1*), Chemokine (C-X-C motif) ligand 2 (*Cxcl2*), Macrophage migration inhibitory factor (*Mif*), Transforming growth factor-β1(*Tgfb1*), Collagen type I alpha (*Col1a*) and Interleukin-10 (*Il10*) are all significantly reduced in mTOR^LEC^ samples. Data are represented as mean + SEM by unpaired two-sided Student’s *t* test, \**p* < 0.05, \*\**p* <0.01, \*\*\**p* <0.001.

### Partial hepatectomy results in necroptosis upon LEC-specific deletion of mTOR

The histopathological analysis of the tissue samples demonstrated the existence of focal necrotic areas exclusively in the livers of mTOR^LEC^ mice after partial liver resection (Figure 7A). As these areas morphologically resembled necroptosis (42, 43), we decided to analyze the protein expression of the established necroptosis marker RIP3K (protein kinase receptor-interacting protein 3) (44). Indeed, RIP3K protein was found to be stabilized in the livers of mTOR^LEC^ mice after partial hepatectomy (Figure 7B), supporting the emergence of necroptosis in this setting. Immunohistochemistry displayed a robust inflammatory response to the focal necrotic areas (Figure 7A), well in line with the established pro-inflammatory consequences of necroptosis (45).

**Figure 7.**
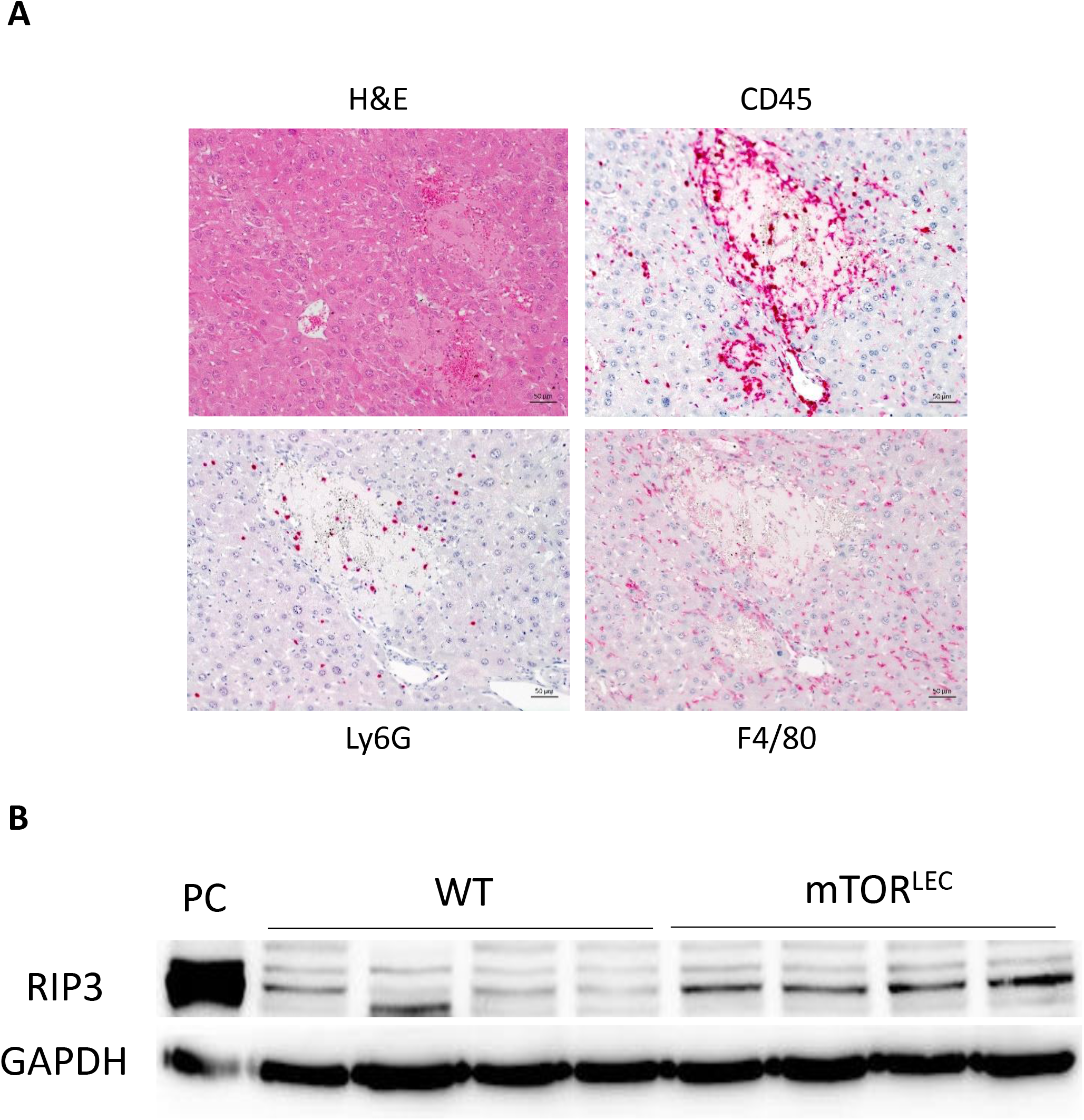
Development of necroptosis in the livers of mTOR^LEC^ mice during liver regeneration. **A**, Morphology of focal necrotic areas in mTOR^LEC^ mice after partial hepatectomy. Sections were stained with H&E as well as antibodies against CD45, F4/80 and Ly6G. **B**, Protein expression of the necroptosis marker RIP3K was analyzed by Immunoblot in whole liver lysates of WT and mTOR^LEC^ mice after partial hepatectomy. A sample from DSS-induced colitis served as the positive control (PC) and GAPDH as the loading control.

**Figure 8.**
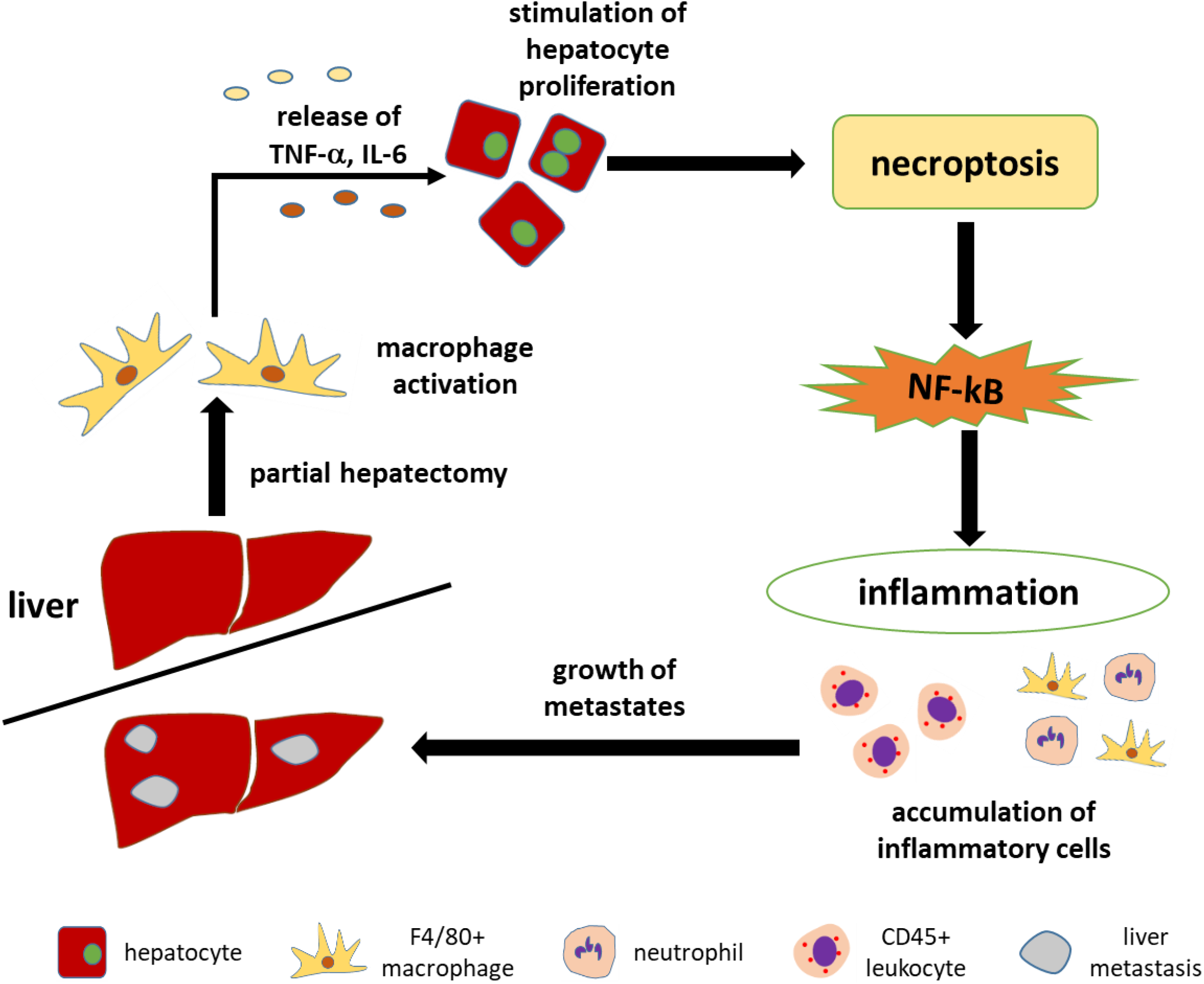
Graphical summary demonstrating the potential sequence of events leading to enhanced metastatic growth after partial liver resection upon LEC-specific mTOR loss.

## Discussion

Our results demonstrate that inhibiting the mTOR kinase in liver epithelial cells aggravates metastatic growth of colon cancer after partial liver resection. We observed elevated inflammatory activity in the livers of mTOR^LEC^ mice after partial hepatectomy and provide experimental evidence to support a functional role of canonical NF-kB in this setting. These results are well in line with clinical experience with mTOR inhibitors as rapamycin and its derivatives are known to induce both systemic and organ-specific (e.g. lung, skin, kidney, intestine) inflammation (46, 47). In addition, LEC-specific loss of mTORC1 (via deletion of the regulatory subunit Raptor) resulted in enhanced hepatic inflammation, characterized by elevated numbers of macrophages as well as T- and B-lymphocytes (48). A functional interdependence of mTOR and NF-kB signaling was reported for the first time by the group of Tony Hunter, noting attenuated NF-kB activity upon activation of mTORC1 (via deletion of the negative mTORC1 regulators TSC1 and TSC2) (40). Shortly thereafter, researchers from Vienna not only confirmed the attenuating function of mTORC1 activation, but went on to show that mTORC1 inhibition (via rapamycin) actually enhanced NF-kB signaling (41). Furthermore, genetic deletion of the mTOR kinase in hematopoietic stem cells resulted in NF-kB activation (49), all well in line with our results. The molecular basis for these findings remained largely elusive until the Superti-Furga lab displayed that mTOR inhibition (via Torin-1 or PP242) markedly suppressed the re-synthesis of IkBα, resulting in enhanced nuclear abundance of p65 and, ultimately, expression of NF-kB target genes upon TNF-α stimulation (50). As TNF-α is of pivotal importance in the early stages of liver regeneration (51), this mechanism could well explain the enhanced NF-kB signaling observed by us. However, the exact molecular nature of the interconnection of mTOR and NF-kB after partial liver resection remains elusive at this point and needs to be addressed by future work.

The association of increased metastasis with elevated canonical NF-kB signaling upon LEC-specific loss of mTOR suggested a functional importance of NF-kB for liver metastasis. This is well in line with published work demonstrating reduced liver metastasis of murine lung cancer and melanoma due to functional inactivation of IKK-β, a kinase of pivotal importance for activation of the canonical NF-kB pathway (52). In this publication, the authors were able to show activated canonical NF-kB signaling in murine livers in response to the establishment of metastases from Lewis lung carcinoma (LLC) cells. Double staining with F4/80 and phospho-IκBα, an established marker of NF-kB activation, demonstrated that the majority of activated NF-kB was found in hepatic macrophages (52). A functional importance of NF-kB in macrophages for metastatic growth was further supported by conditional gene targeting of IKK-β. Of note, LEC-specific deletion of IKK-β had no impact on the metastatic load in this publication. The obvious conflict with our results is most likely explained by the different experimental setups. We only observed activation of the canonical NF-kB pathway in hepatocytes under conditions of mTOR deficiency and liver regeneration. Taken together, our data suggest that under certain conditions the canonical NF-kB pathway in hepatocytes is indeed able to impact on the progression of liver metastases.

The PI3K/Akt/mTOR signaling pathway is activated by various growth factors and mitogens during the process of liver regeneration (53). Successful and timely completion of liver regeneration after liver surgery is of paramount importance for postoperative rehabilitation and the overall prognosis of the patient. Independent research groups were able to show that treatment with rapamycin and rapalogs did not significantly delay liver regeneration in different rodent models of partial hepatectomy (54–56). Of note, the systemic administration of mTORC1 inhibitors was not without effect as it significantly reduced hepatocyte proliferation, enhanced apoptosis of hepatic cells and reduced expression of growth factors at different time points in the first week after partial hepatectomy (54–56). However, this neither affected the timely completion of the regenerative process nor the survival of the animals in a significant way. In principle, these results were confirmed in a mouse model displaying defective mTORC1 specifically in liver epithelial cells (48). Again, mTORC1 deficiency resulted in early defects of hepatocyte mitosis and enhanced hepatocellular damage early after partial liver resection, but overall liver regeneration and animal survival was not affected. While these results argue for a compensable role of mTORC1 for liver regeneration, a different picture emerges when mTORC2 signaling is analysed. Mice harbouring a liver epithelial cell-specific inactivation of mTORC2 (due to Cre/loxP-mediated deletion of *Rictor*) showed higher mortality after partial hepatectomy, potentially due to reduced activation of Akt (57). These results suggest that the two basic components of mTOR encompass different functional roles for successful liver regeneration after partial hepatectomy. Our results are seemingly not in line with this suggestion as we did not observe defective liver regeneration upon LEC-specific inactivation of the mTOR kinase (resulting in loss of both mTORC1 and mTORC2). Of note, our mouse model is characterized by a rather low knock-out efficiency, much lower compared to the efficiency reported for Albumin-Cre-driven deletion of *Raptor* and *Rictor*, respectively (48, 58). This points to a paramount importance of mTOR signaling for the function and survival of hepatocytes and bile duct epithelial cells. It can be concluded that the low knock-out efficiency observed in our mice explains the inconsistency with the work of Xu *et al.* (57), who have noted a significant contribution of mTORC2 in liver epithelial cells for regeneration after partial hepatectomy.

Necroptosis is a form of programmed cell death and a characteristic response of caspase-8-deficient cells to TNF-α or other stimulators of the extrinsic pathway of triggering apoptosis (42, 59, 60). We observed necroptosis in LEC-specific mTOR KO mice after partial hepatectomy in a situation of enhanced activity of the canonical NF-kB pathway. This conflicts with reports demonstrating necroptosis in mice with robustly reduced NF-kB activity in liver epithelial cells due to deletion of IKKα/β or TAK-1 (TGF-β-activated kinase 1) (43, 61). Of note, our experimental setup did not permit to unravel the exact functional connection between necroptosis and enhanced NF-kB activity. While the results mentioned above support a necroptosis-inhibiting role for NF-kB reminiscent of the well-established anti-apoptotic function of NF-kB, our observations could be explained by an activation of the canonical NF-**k**B pathway downstream of necroptosis. In fact, this is supported by published data from the group of Junying Yuan (who coined the term “necroptosis” in 2005), demonstrating robust activation of the canonical NF-kB pathway in various cell lines in response to activation of necroptosis (62). What’s more, the Yuan group reported enhanced expression and secretion of pro-inflammatory cytokines as a consequence of necroptosis-induced NF-kB activation, a finding well in line with our results. Taken together, this suggested that the LEC-specific inactivation of mTOR results in the activation of hepatocellular necroptosis in the context of liver regeneration due to partial hepatectomy. However, this is not supported by the published literature, where several independent groups have reported a functional importance of mTOR activation for the execution of necroptosis (63–65). We cannot rule out the possibility that the focal necrotic areas in the mTOR^LEC^ mice originated from mTOR-proficient hepatocytes. It is at least conceivable that the intercellular communication in the liver is modified by the presence of mTOR-deficient hepatocytes in such a way that a regenerative stimulus results in focal necroptosis. While this seems to be a rather daring hypothesis, cell-intrinsic effects of the mTOR loss might be better suited to explain our observation. Specifically, a potential impact of the loss of mTOR on the activity of caspase-8 seems mandatory to address.

## Author Contributions

Conceptualization, L.J., R.E., M.S., U.P.N. and T.C.; Experimentation L.J., R.E., S.J., J.R., A.K., M.E., H.D., C.L-I., L.R.H.; Data Analysis and Visualization, L.J., R.E., S.J., A.K., M.E., H.D., C.L-I., D.M., L.R.H. and T.C.; Writing, L.J., R.E. and T.C.; Supervision, Resources, and Funding Acquisition, U.P.N. and T.C.; Project Administration, T.C.

## Acknowledgements

Research in the Cramer lab was supported by grants from the Deutsche Forschungsgemeinschaft (CR 133/2-1 until 2-4 and CR 133/3-1). L.J. was supported by the China Scholarship Council (CSC, file NO. 201706260268). The excellent technical assistance of Nelli Neuberger and Ilka Sauer is highly appreciated. We are grateful to Pavel Strnad (Gastroenterology, RWTH Aachen University Hospital) for helpful discussions and for providing control reagents. This work has partly been presented as a short talk at the 2019 United European Gastroenterology Week in Barcelona, Spain.

## Supplementary Figure Legends

**Figure S1. A**, Mtor PCR on genomic DNA extracted from isolated hepatocytes of WT and mTOR^LEC^ mice. **B**, Western blot analysis of isolated hepatocytes and liver whole protein between WT and mTOR^LEC^ mice. **C**, Murine hepatocytes were stimulated with insulin (10 nmol/l, 30 min), mTOR, phospho-p70S6K and phospho-AKT ser473 expression were analyzed via Western Blot, β-actin served as the loading control.

**Figure S2. A**, The Pearson correlation of KO versus WT liver proteomes in scatter plots. **B**, Volcano plot featuring liver proteome data, with the mean difference of LFQ intensities between mTOR^LEC^ and WT tissues on the × axis versus statistical significance on the y axis (−log10 of the p value). Significantly regulated proteins are shown in red (down in mTOR^LEC^) and green (up in mTOR^LEC^).

**Figure S3.** STRING PPI (protein–protein interaction) network and pathway analysis of proteins with downregulation at least one standard deviations from the log2 transformed median values in mTOR^LEC^/WT liver tissue. Proteins which belong to the GO (Gene Ontology) pathway “RNA binding” are highlighted in red and those which belong to the KEGG (Kyoto Encyclopedia of Genes and Genomes) pathway “ribosome” are highlighted in blue. Unconnected nodes were removed.

**Figure S4.** Effect of LEC-specific mTOR deletion on proliferation and apoptosis of hepatocytes. Immunohistochemistry against phosphorylated Histone H3 and activated caspase-3 was performed. Positive cells were quantified with ImageJ and results are shown as bar graphs.

**Figure S5.** Analysis of autophagy in WT and mTOR^LEC^ livers. **A**, Western blot against p62 and LC3B was performed on whole protein samples from primary murine hepatocytes, β-actin served as the loading control. Densitometry was used to quantify protein expression relative to the loading control. **B**, Electron microscopy (EM) was performed on liver samples from WT and mTOR^LEC^ mice, arrows demonstrate autophagosomes.

**Figure S6.** Proliferation of tumor cells and hepatocytes adjacent to metastases was analyzed via immunohistochemistry against phosphorylated Histone H3.

**Figure S7.** Schematic diagram of the applied murine models of colon cancer (MC38 cells) liver metastases. **A**, Mirroring the initial metastatic phase and **B**, representing the situation after surgical resection of liver metastases.

**Figure S8.** Proliferation of tumor cells and hepatocytes adjacent to metastases after partial hepatectomy was analyzed via immunohistochemistry against phosphorylated Histone H3.

