## Supplementary figures and images for "Enhanced inflammatory response mediated by parenchymal cells associates with resistance towards mTOR inhibition"

### Supplemental Figures

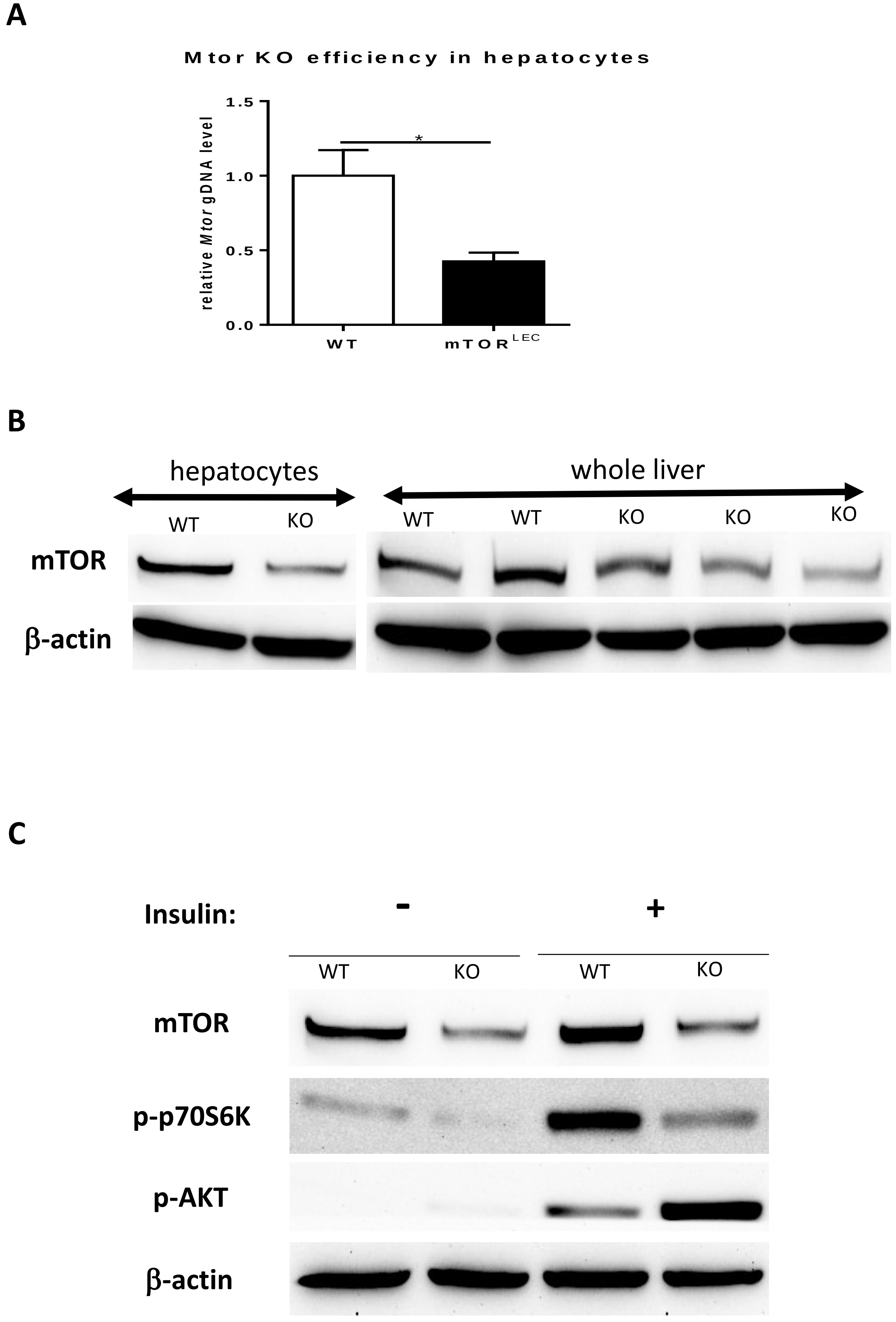

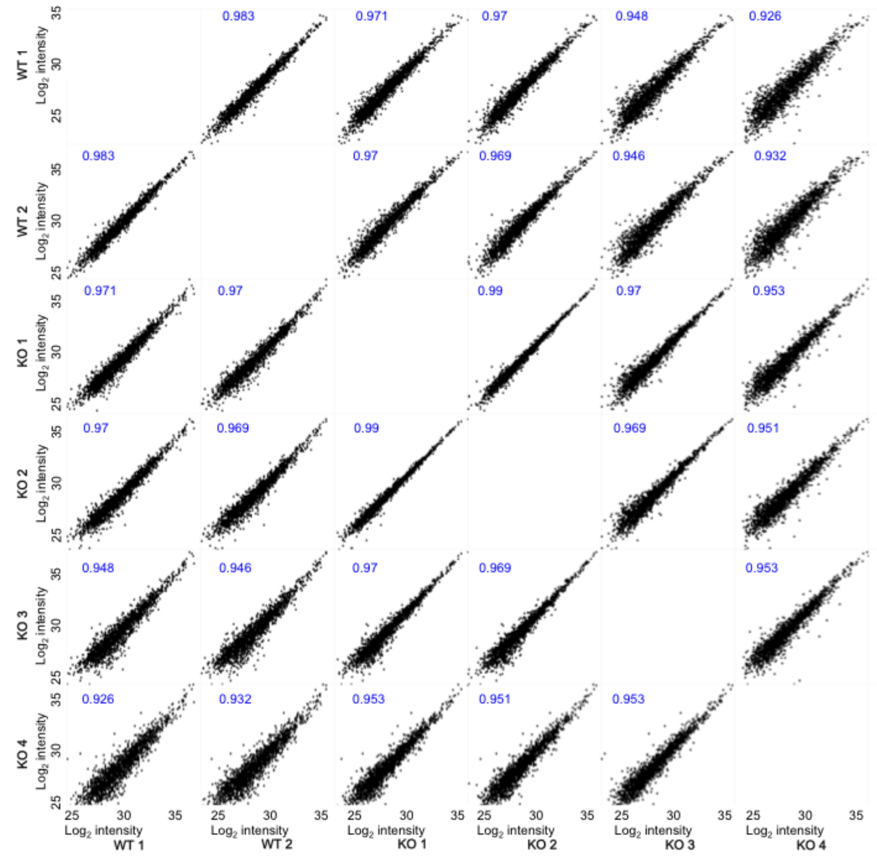

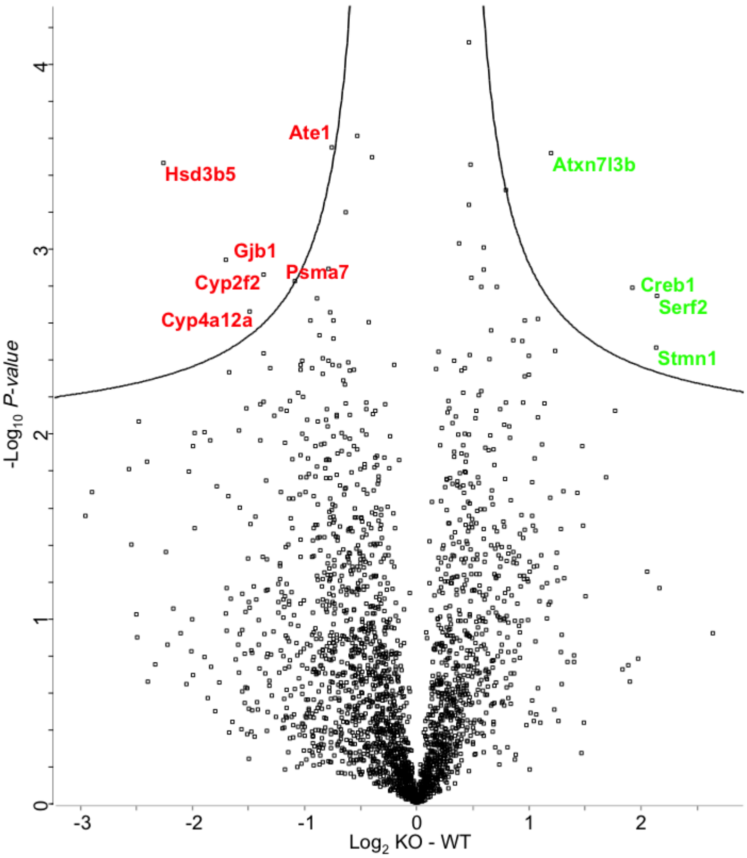

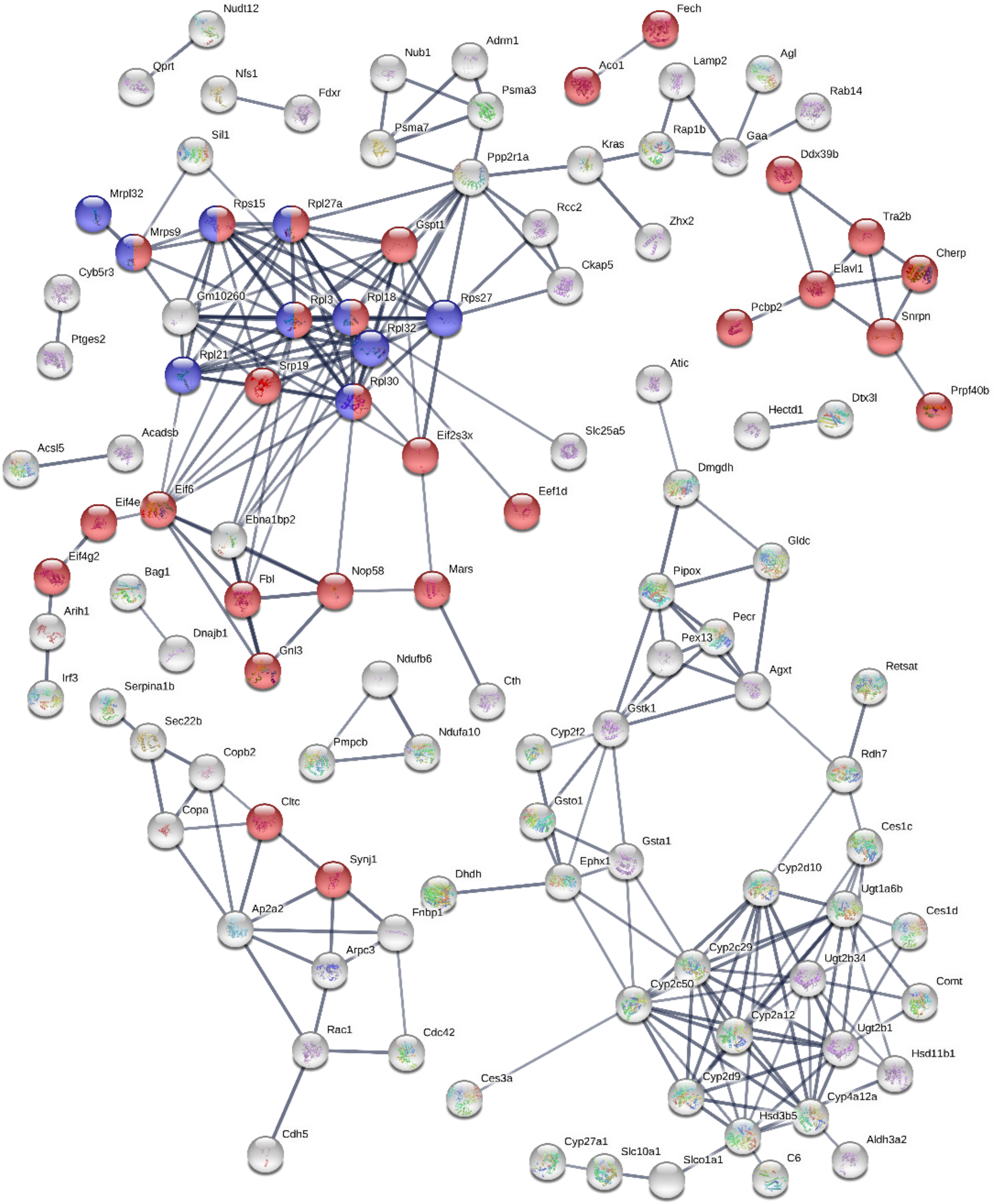

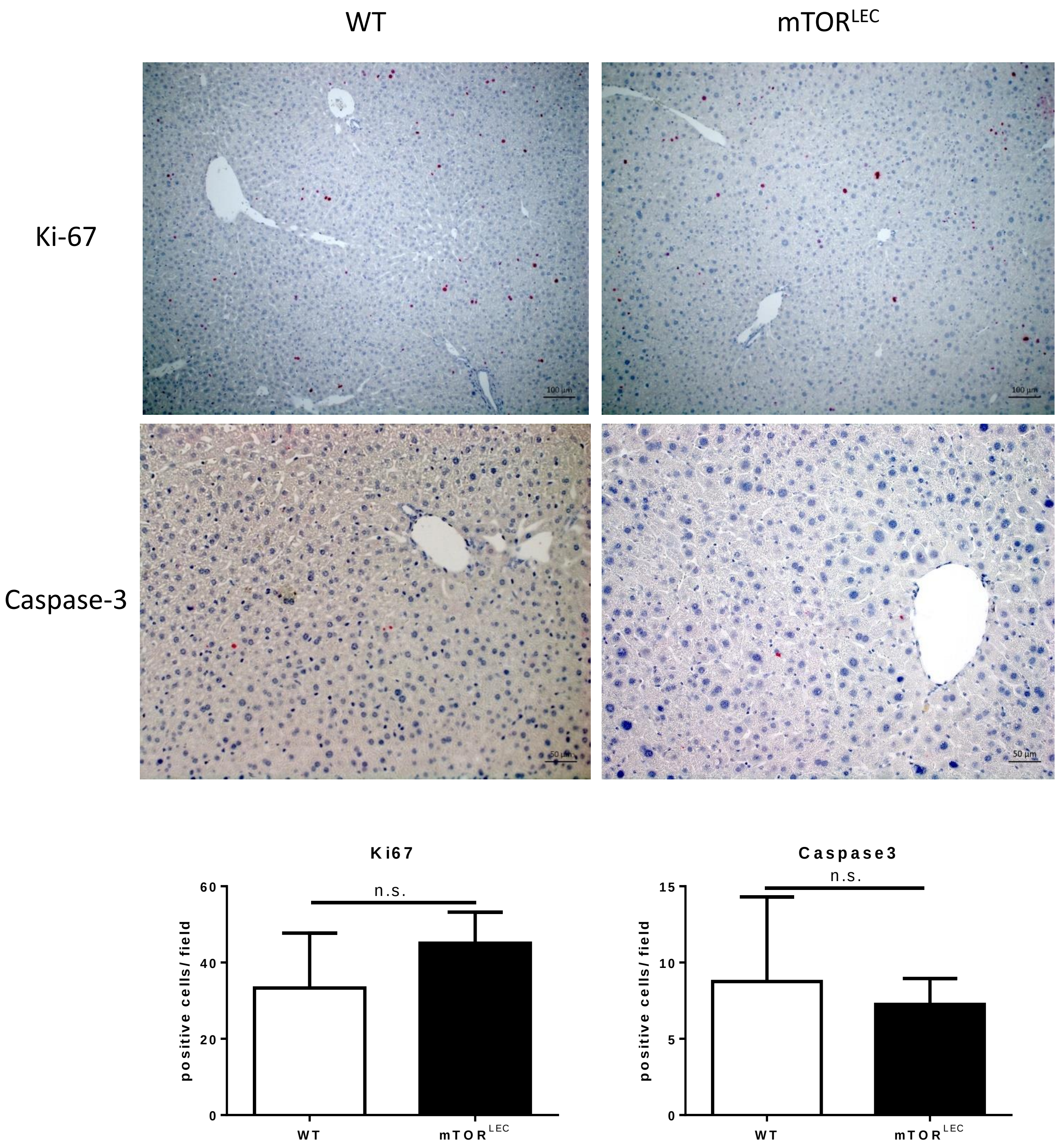

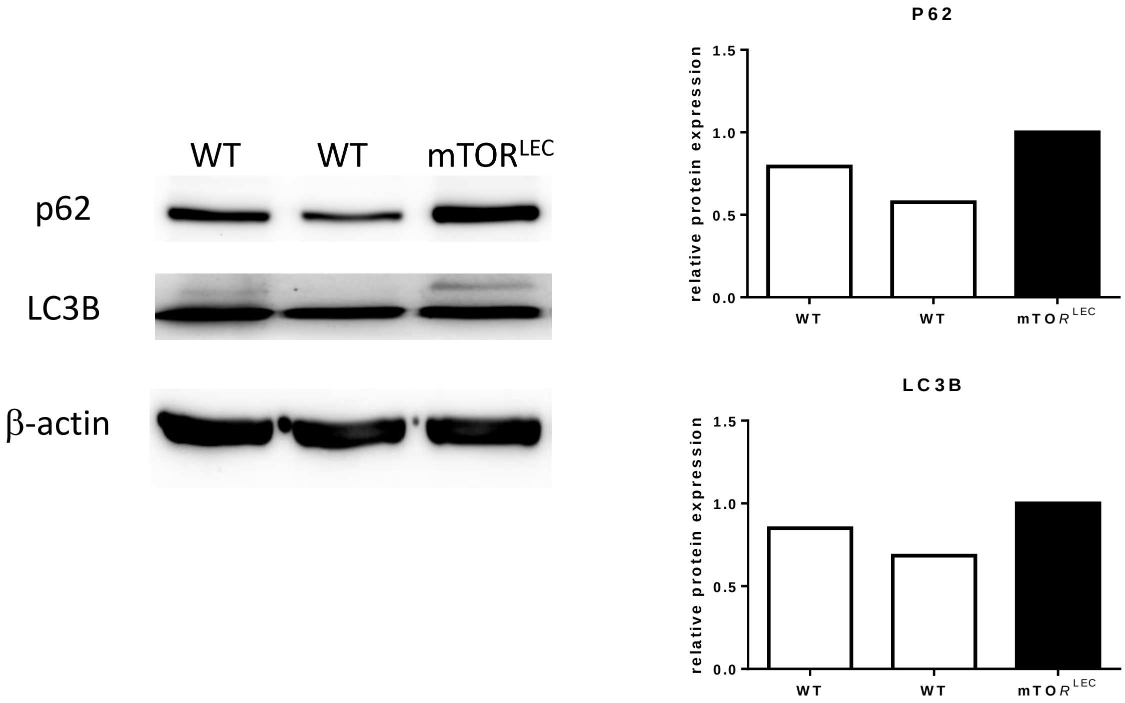

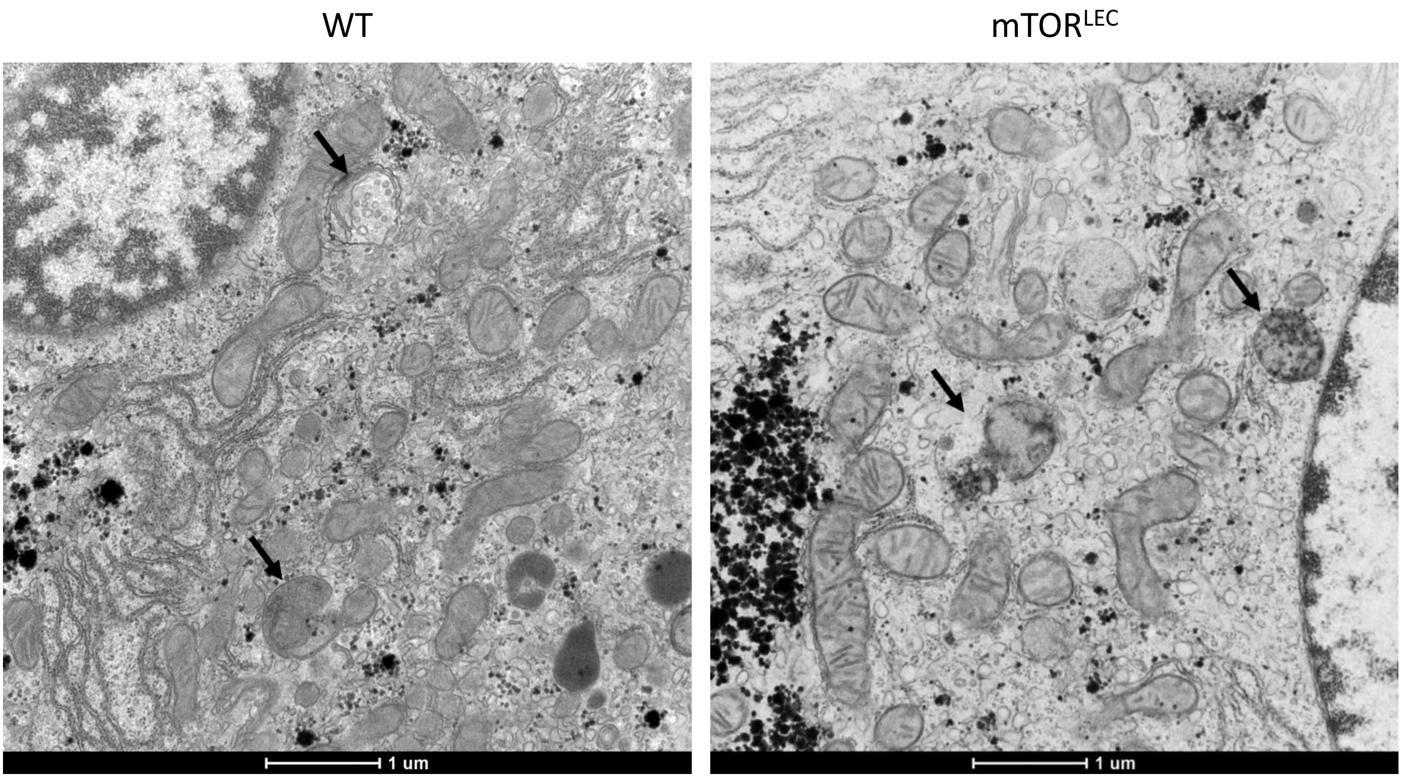

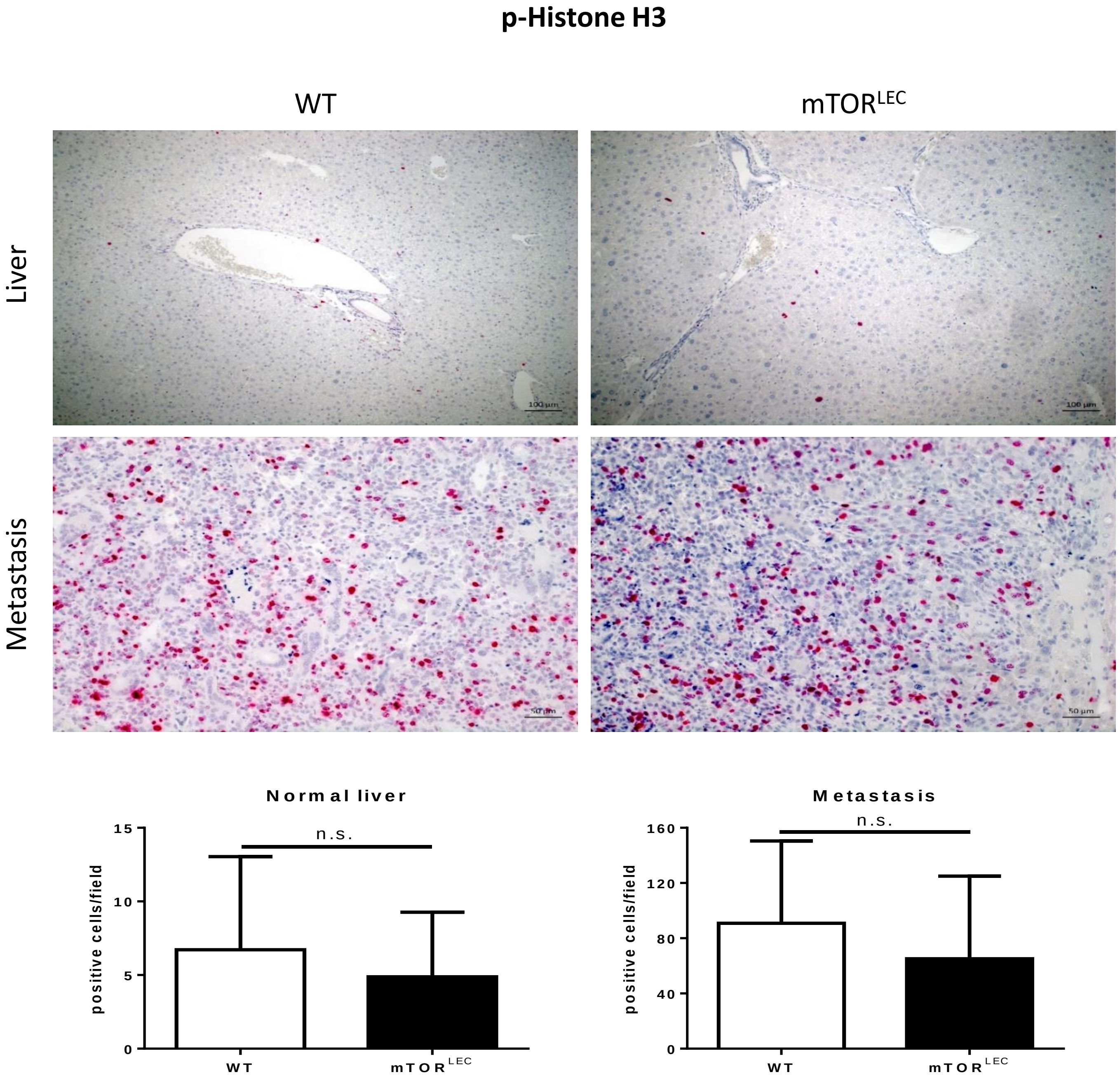

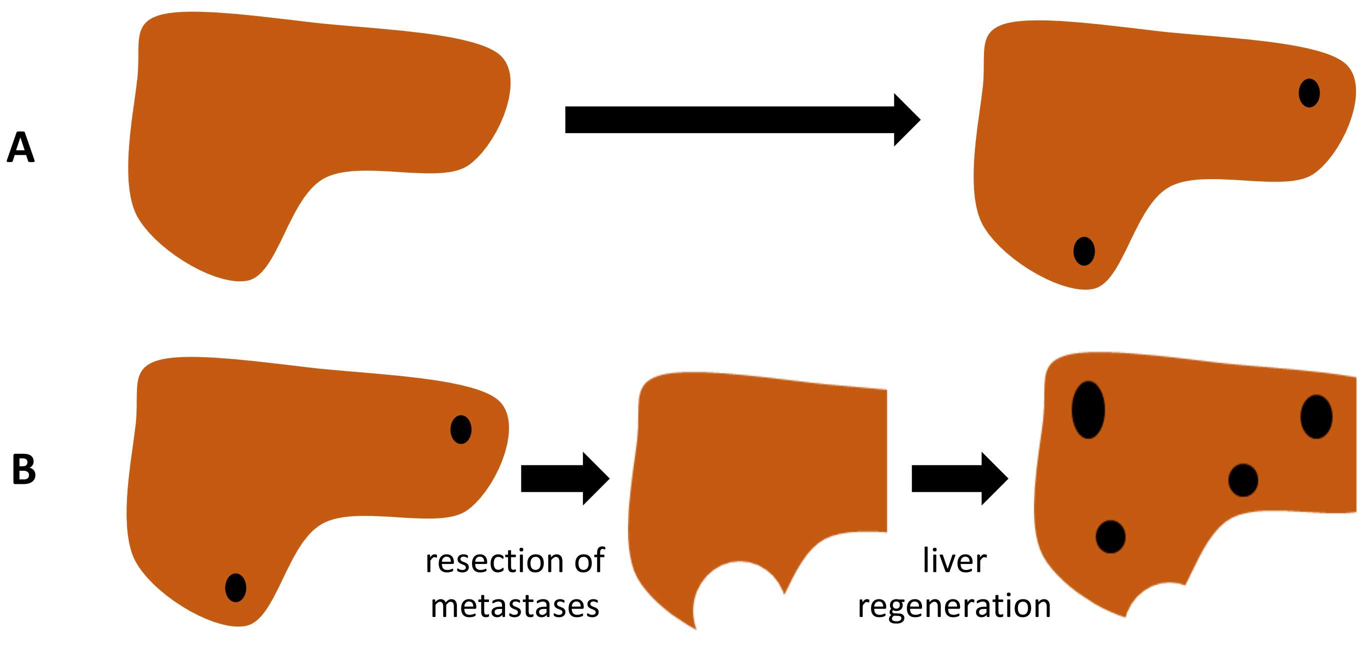

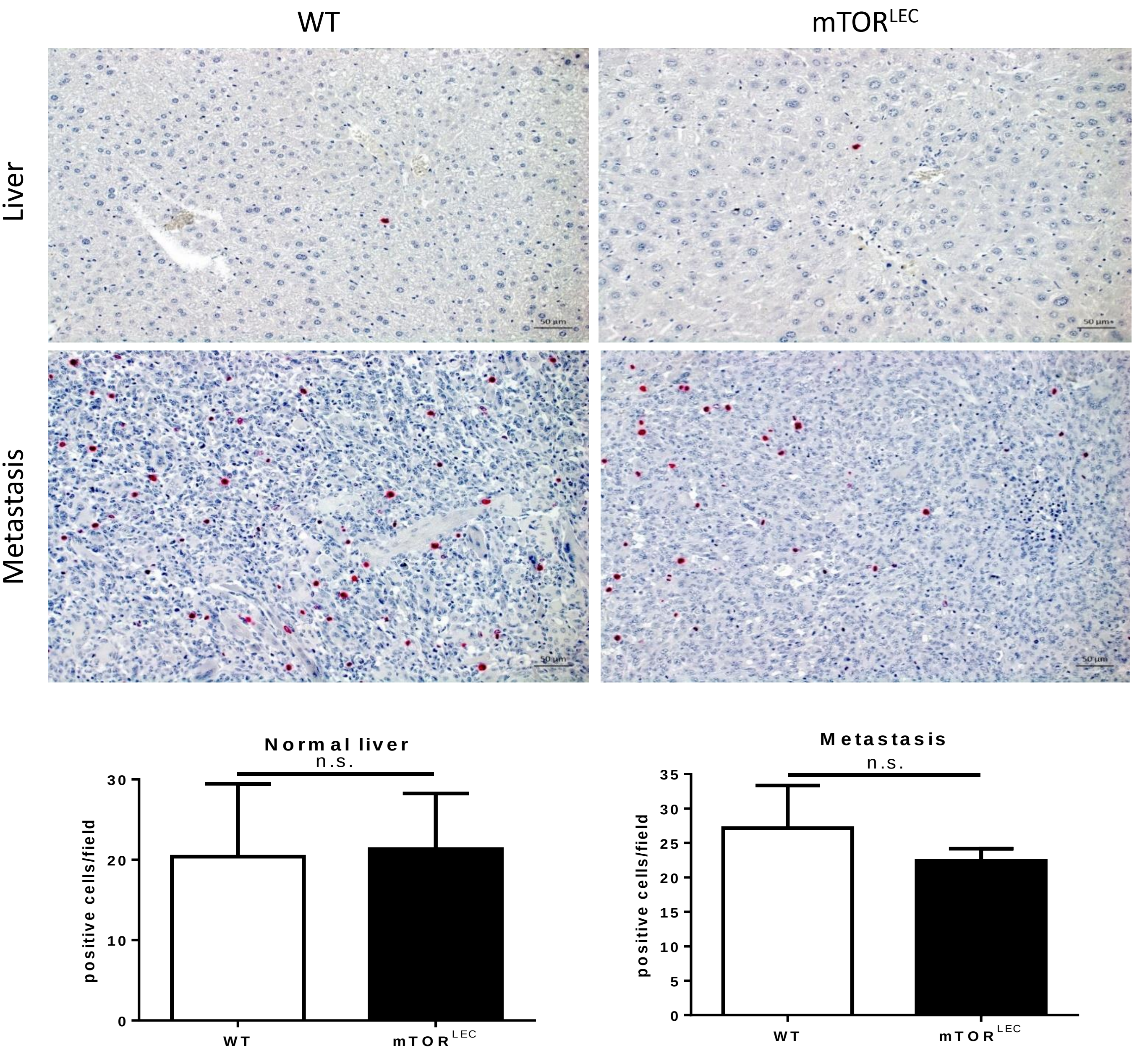
